# Fluorescent Chiral Quantum Dots to Unveil Origin-Dependent Exosome Uptake and Cargo Release

**DOI:** 10.1101/2023.12.20.572689

**Authors:** Gaeun Kim, Runyao Zhu, Youwen Zhang, Hyunsu Jeon, Yichun Wang

## Abstract

Exosomes are promising nanocarriers for drug delivery. Yet, it is challenging to apply exosomes in clinical use due to the limited understanding of their physiological functions. While cellular uptake of exosomes is generally known through endocytosis and/or membrane fusion, the mechanisms of origin-dependent cellular uptake and subsequent cargo release of exosomes into recipient cells are still unclear. Herein, we investigated the intricate mechanisms of exosome entry into recipient cells and the intracellular cargo release. In this study, we utilized chiral graphene quantum dots (GQDs) as representatives of exosomal cargo, taking advantage of the superior permeability of chiral GQDs into lipid membranes, as well as their excellent optical properties for tracking analysis. We observed a higher uptake rate of exosomes in their parental recipient cells. However, these exosomes were predominantly entrapped in lysosomes through endocytosis (intraspecies endocytic uptake). On the other hand, in non-parental recipient cells, exosomes exhibited a greater inclination for cellular uptake through membrane fusion, followed by direct cargo release into the cytosol (cross-species direct fusion uptake). We revealed the underlying mechanisms involved in the cellular uptake and the subsequent cargo release of exosomes depending on their cell-of-origin and recipient cell types. This study envisions valuable insights into further advancements in the effective drug delivery using exosomes, as well as a comprehensive understanding of cellular communication, including disease pathogenesis.

## 1. INTRODUCTION

Exosomes are a subset of extracellular vesicles, with diameters between 50 and 150 nm, secreted naturally by most eukaryotic cells.^1, 2^ These biological nanovesicles contain various biomolecule cargo, such as nucleic acids, peptides, proteins, and metabolites, originating from their parent cells.^3, 4^ Recently, exosomes have come into the spotlight due to their functions in intercellular communication and their ability to deliver biomolecule cargo.^5^ They are being extensively investigated as drug delivery carriers, attributed to their intrinsic characteristics, such as nanoscale size, biocompatibility, low toxicity, low immunogenicity, targeting to specific cells, and the potential to cross biological barriers.^6^ These exceptional properties highlight exosomes as promising nanocarriers for drug delivery system to treat a wide range of human diseases, such as cancers, neurodegenerative diseases, and cardiomyopathies.^7, 8^

Despite the potential benefits, the utilization of exosomes in clinical application remains challenging. One of the main challenges is the inefficient specific targeting and cellular uptake.^9^ Hence to design and develop therapeutic exosomes for drug delivery, in-depth comprehension of mechanisms for cellular uptake and the intracellular cargo release is essential. For instance, the cellular entry mechanism is crucial for optimizing drug delivery strategies to enhance therapeutic efficacy,^10^ while deciphering cargo release mechanisms is imperative for designing precise cargo delivery and release to intracellular targets efficiently.^11^ Exosomes have a tendency to target their cell-of-origin, resulting preferential uptake into their parental recipient cells compared to exosomes from different origins.^12–16^ This trend has been demonstrate both *in vitro* and *in vivo*.^17^ Upon reaching the recipient cells, exosomes undergo through two possible uptake pathways: 1) endocytosis that is mediated by the interaction of ligands on exosome membranes with specific receptors on cell membranes, and 2) direct fusion with the plasma cellular membrane.^18, 19^ The specific mechanism underlying the cellular uptake of exosomes to deliver their cargo largely depends on how recipient cells interact with exosomes, however, this detail remains largely unexplored.^12, 19^ Therefore, further investigations are necessary to unravel mechanisms governing the two pathways of cellular uptake for exosomes, as well as the following intracellular cargo release.

To investigate the cellular uptake and subsequential cargo release of exosomes, devising a comprehensive tracking methodology is of utmost importance. Conventional membrane staining dyes, like Carbocyanine dyes^20^ and Styryl dyes^21^, have been used widely in biological studies for exosome imaging tracking through membrane staining or labeling. However, these dyes might lead to inaccurate assessments due to uneven staining of lipids under non-optimal and inconsistent conditions.^22^ The non-specific lipid staining potentially cause false-positive results due to the polarity of solvent used for dye dissolution.^23^ Furthermore, by binding to surface molecules or incorporating with the plasma membranes, lipid-like or lipophilic molecules, such as Nile Red, might induce significant changes in cellular membrane properties, potentially triggering signal transduction within the cell and altering the fluidity of membrane.^24, 25^ In particular, lipophilic dye tends to aggregate in salt-containing solutions such as phosphate-buffered saline (PBS), which is commonly used to suspend exosomes. The aggregation of these dye molecules hinders their efficient integration into vesicle membranes and leads to non-specific fluorescence signals.^26, 27^ Therefore, depending on the type and concentration of the staining dye used, it has the potential to influence the properties of cell membranes, cellular processes, or interactions with other cells or molecules, which leads to erroneous results.^28^ Moreover, not only labeling exosomes but also to further explore their journey from extracellular interactions to intracellular cargo release, the meticulous cargo tracking approach for in-depth analysis is essential for this study.^29^ Several studies reported techniques of loading the probes for exosomal cargo tracking through endogenous approaches such as isolation of exosomes from cells that were genetically modified to express reporter proteins^30^ and exogenous approaches such as loading fluorescent-tagged proteins into exosomes through process of membrane rupture and revesiculation.^31^ However, a low yield of exosomes containing biologics remains a challenge when utilizing passive endogeneous loading methods, which usually yield below 30% due to the intricate process of exosome biogenesis and cargo sorting mechanisms.^32^ In addition, most active exogeneous loading methods potentially damage lipids and denature proteins, which could lead to the degradation of the intrinsic exosomal structure and alteration of their biological functions.^33^ Hence, an approach that loads sufficient probes representing cargo of exosomes without damaging the integrity of their structure is essential for investigating the cell-exosome communication profile.

In this study, we utilized fluorescent chiral graphene quantum dots (GQDs) as a representative cargo of exosomes to investigate the differential cellular uptake and cargo release of exosomes depending on their cell-of-origin and recipient cell types. These chiral GQDs were derived by surface functionalization with *D*-cysteine (*D*-GQDs), exhibiting high permeability into exosomes through the nanoscale chirality and its interaction with lipid membranes.^34, 35^ More importantly, the size and morphology of exosomes remained intact after the successful loading of *D*-GQDs.^35^ In addition, chiral GQDs exhibit unique optical properties, particularly their fluorescence emission, allowing for their application in bioimaging and biosensing.^36^ Analyzing GQD accumulation through fluorescence-based techniques provides valuable information to estimate their uptake *in vitro* and a better understanding of their distribution *in vivo*.^36, 37^ By using confocal laser scanning microscopy (CLSM) imaging analysis, mass spectrometry (MS)-based proteomics, Western blot, and fluorometry, we revealed that exosomes derived from the same cell-of-origin (intraspecies exosomes) are preferentially taken up by their parental recipient cells through receptor-ligand interaction-mediated endocytosis. The cellular uptake of intraspecies exosomes is 1.4∼3.2 times higher compared to exosomes derived from different cell-of-origin (cross-species exosomes). On the other hand, cross-species exosomes were predominantly taken up by non-parental recipient cells through direct membrane fusion, with a strong correlation value (PCC>0.7). In contrast, intraspecies exosomes exhibited a lower correlation value of 0.3∼0.5 with parental recipient cells. We revealed that intraspecies exosomes were more inclined to interact with receptors on their parental recipient cells and were taken up through endocytosis, followed by the entrapment of exosomal cargo within lysosomes. In contrast, cross-species exosomes tended to fuse directly with the membranes of non-parental recipient cells, releasing their cargo into the cytosol. Overall, our study offers a valuable understanding of the distinct mechanisms of exosome uptake and cargo release that were not fully comprehended before.

## 2. MATERIALS AND METHODS

### 2.1. Materials

Carbon nanofibers (719803), and PKH26 (MINI26-1KT) were purchased from Sigma-Aldrich (MO, USA). Sulfuric acid (BDH3068-500MLP), Nitric acid (BDH3044-500MLPC), and Fetal bovine serum (FBS; 1300-500) were purchased from VWR (PA, USA). Dialysis membrane tubing (MWCO: 1 kD; 20060186) was purchased from Spectrum Chemical Manufacturing Company (NJ, USA). 1-ethyl-3-(3-dimethyl-aminopropyl) carbodiimide (EDC; 22980), VybrantTM Multicolor Cell-Labeling Kit (DiO, DiI; V22889), and GibcoTM Trypsin-EDTA (25200072) were purchased from Thermofisher Scientific (MA, USA). N-hydroxysuccinimide sodium salt (Sulfo-NHS; 56485), and *D*-cysteine (A110205-011) were purchased from AmBeed (IL, USA). Nitrocellulose membrane (1662807), and Clarity Max Western Enhanced Chemiluminescence (ECL) Substrate (1705060) were purchased from Bio-Rad (CA, USA). RIPA Buffer (9806), anti-mouse Horseradish Peroxidase (HRP)-linked secondary antibody (7076), and anti-rabbit HRP-linked secondary antibody (7074) were purchased from Cell Signaling Technology (MA, USA). Anti-CD9 antibody, anti-CD63 antibody, anti-CD81 antibody, and anti-rabbit HRP-linked secondary antibody (EXOAB-KIT-1) were purchased from System Biosciences (CA, USA). Anti-beta-actin antibody (sc-47778), anti-neuregulin1 antibody (sc-393006), and anti-GalNAc antibody (sc-393370) were purchased from Santa Cruz Biotechnology (TX, USA). Anti-TGF-beta1 antibody (21898-1-AP) was purchased from Proteintech (IL, USA). LysoViewTM 594 (70084), Hoechst 33342 (40046), and CellBrite Green Cytoplasmic Membrane Dye (30021) were purchased from Biotium (CA, USA). CytoTraceTM Red CMTPX (22015) was purchased from AAT Bioquest (CA, USA). Minimum essential medium eagle (MEM; 10-010-CV), and Phosphate-buffered saline (PBS; 21-040-CM) were purchased from Corning (NY, USA). Antibiotic antimycotic (15240096) was purchased from Fisher Scientific (MA, USA). 4% Paraformaldehyde (PFA; 15735-50S), and UranyLess (22409) were purchased from Electron Microscopy Sciences (PA, USA). Tween-20 (BTNM-0080) was purchased from G-Biosciences (MO, USA).

### 2.2. Instruments

Transmission electron microscopy (TEM; JEOL 2011, JEOL Ltd., Tokyo, Japan) was used to confirm the nanostructures. Zeta potential values were characterized by Zetasizer (Malvern Zetasizer Nano ZS, Malvern Instrument Ltd., Worcestershire, UK). Nanoparticle Tracking Analysis (NTA; NanoSight LM10 system, Malvern Instrument Ltd., Worcestershire, UK) was used to measure the size distribution and concentration of nanoparticles. Confocal Laser Scanning Microscopy (CLSM; A1R-MP Laser Scanning Confocal Microscopy, Nikon, Tokyo, Japan) was operated to observe the cellular uptake and penetration of chiral GQDs into exosomes. Fluorescence was measured by a plate reader (Infinite 200 PRO, Tecan, Männedorf, Switzerland). Circular dichroism (CD) spectrometer (Jasco J-1700 Spectrometer, Jasco International Company, MD, USA) was used to measure the absorbance of polarized light were measured. Mass spectrometer (MS; Thermo Q-Exactive HF, Thermo Fisher, MA, USA) was used to identify the proteins. Western blot images for protein immunodetection were acquired using a bioimaging system (C400 Bioanalytical Imager, Azure Biosystems, CA, USA). Humidified incubator (MCO-15AC, Osaka, Japan) was used to maintain and incubate all cell lines used in this study.

### 2.3. GQD Synthesis and Chiral Functionalization

The GQDs were synthesized using a modified protocol based on our previous report ^34, 35^ as follows: A 40 mL mixture of sulfuric acid and nitric acid (3:1, v/v) and 0.4 g of carbon nanofibers were sonicated for 2 h. After being mechanically stirred for 6 h at room temperature, the mixture was heated to 120 °C and allowed to react continuously for 10 h. Then, the mixture was cooled, diluted with ice-cold deionized water, and pH was adjusted to 8 by adding sodium hydroxide. After 3 d of dialysis for purification, we got GQDs with the final concentration of 1 mg/mL. To functionalize the GQDs with chirality, the carboxylic group of GQDs was connected to the amine group of *D*-cysteine by EDC/NHS coupling reaction ^35, 38^ as follows: 20 μL of EDC (100 mM) was added to 2 mL of GQDs (100 μM) solution, followed by 10 min of stirring. Then, 40 μL of Sulfo-NHS (100 mM) was added and sonicated for 40 min under the ice-water bath. In ordre to remove excess EDC and Sulfo-NHS, the mixture was filtered with a 1 kDa centrifuge tube and rinsed three times. Finally, 40 μL of *D*-cysteine (100 mM) was added and the mixture was stirred for 2 h. The excess *D*-cysteine was removed by the dialysis membrane. The chiroptical activity was measured using a CD spectrometer, and the fluorescence property was measured using a plate reader. The morphology of chiral GQDs was observed using TEM. The SigmaPlot 10.0 (Systat Software Inc., CA, USA) was used to generate plots of the analyzed data.

### 2.4. Cell Culture

Human hepatoma cells (HepG2), mouse fibroblast cells (3T3), and human cervical adenocarcinoma cells (HeLa) were cultured in MEM supplemented with 10% FBS and 1% Antibiotic-Antimyotic in a humidified incubator at 37°C with 5% CO_2_. Cells were washed with 1X PBS, trypsinized with trypsin-EDTA for passages, and incubated at least 1 d before any experiment. Cells were stained with several dyes for imaging under CLSM: CytoTrace Red CMTPX dye for cytosol area staining; CellBrite Green Cytoplasmic Membrane dye and DiO dye for membrane staining; LysoView 594 dye for lysosome staining; Hoechst 33342 dye for nuclei staining. Cells were fixed with 4% PFA for 15 min before imaging.

### 2.5. Isolation and Characterization of Exosomes

Once HepG2, 3T3, and HeLa cells covered 70-80% of the flask, cell-culture medium (CM; MEM with 10% FBS and 1% Antibiotic-Antimyotic) was changed into serum-free CM after washing three times with 1X PBS. The serum-free CM was collected after 48 h. Then, the serum-free CM was filtered with Pore Size 0.22 µm Vacuum Filtration Systems (10040-460; VWR, PA, USA) to remove undesired large debris. The filtered CM was filtered through 0.05 µm pore size hydrophilic membranes (111103; Cytiva, MA, USA) under mild-vacuum filtration conditions for 2 d. Then, the CM was washed with 1X PBS buffer and concentrated in 100 kDa ultrafiltration centrifugal devices (Spin-X UF Concentrator; Corning, NY, USA). With the negative staining by UranyLess, the morphology and nanostructure of exosomes were observed with TEM and the size distribution of 55-65 exosomes was analyzed using ImageJ software. NTA measurements were performed using diluted exosomes (200 times dilution; 5 μL in 1 mL of PBS buffer) and analyzed with NTA 3.3 analytical software suite. Surface charge of exosomes was measured by a zetasizer. To activate the disposable zeta potential cuvette (DTS1070, Malvern Instrument Ltd., Worcestershire, UK), it was rinsed once with 100% ethanol and washed twice with distilled water. Exosomes diluted into 2-3 × 10^6^ exo/mL were loaded in the cuvette and 5 measurements per sample were made with 30-100 autoruns for each using the Hückel model. Exosomal marker expressions (CD9, CD63, and CD81) were validated through western blot. Exosomal membranes were stained with PKH26 and DiI dye for imaging with CLSM. All characterization procedures were performed at room temperature. The SigmaPlot 10.0 was used to generate plots of the analyzed data.

### 2.6. Permeation of Chiral GQDs into Exosomes

*D*-GQDs concentrations of 12 and 15 μM were incubated with 1×10^9^ particles/mL exosomes at room temperature with protection from light. The incubated mixture was then washed with 1X PBS buffer in a 100 kDa cellulose centrifugal filter 4 times to remove excessive *D*-GQDs. The permeation efficiency of *D*-GQDs into exosomes (*D-*GQDs/exo) was indirectly determined by statistical analysis with a formula from our previous report:^35^

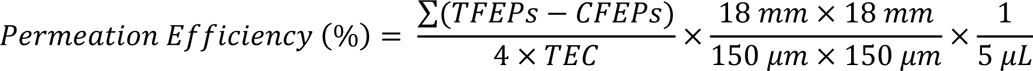

5 μL of *D*-GQDs/exo loaded on the Poly-L-Lysine coated slides (63410; Electron Microscopy Sciences, PA, USA) and was fully covered with 18 mm × 18 mm square cover glasses (2845-18; Corning, NY, USA). The covered area was scanned under a 100× objective with a normal 150 μm × 150 μm field size. Each scan contained around 25 z-stack images (step size of 0.13 μm) of appearing/disappearing blue fluorescent dots. Four z-stack images were captured through random domains. The z-stack images were set manually to adjust the thresholds and match the diameter size range of exosomes (40-140 nm). Then, by counting the blue-fluorescent lit-up *D*-GQDs/exo (due to permeation) using ImageJ software, we can get the total fluorescent exosome particles (TFEPs). Colocalized fluorescent exosome particles (CFEPs) of z-stack images, between successive image sets, were counted via ImageJ software with JACOPx Plugin. The total exosome concentration (TEC) was measured by NTA in particles/mL. For additional validation, the permeated *D*-GQDs were quantified by fluorometric way. The assessment of fluorescent recovery was conducted before and after lysis of exosomes with biocompatible surfactants, Tween-20. The samples were loaded into a 96-well black plate (3991; Corning, NY, USA), and fluorescence intensity (FI) was measured using a plate reader with excitation at 360 nm and emission at 510 nm. The SigmaPlot 10.0 was used to generate plots of the analyzed data.

### 2.7. Cellular Uptake Imaging of Exosomes

Cellular uptake images were obtained by CLSM. All cells were washed out with 1X PBS and fixed with 4% PFA before imaging. To observe the cellular uptake of exosomes, *in vitro* experiments were conducted as follows: CytoTraceTM Red CMTPX pre-stained cells were cultured at 5 × 10^4^ cells per well in 8-well confocal plate (Lab-Tek II Chambered Coverglass, 155409; Thermofisher Scientific, MA, USA) and incubated overnight at 37°C under 5% CO_2_. Three different types of *D*-GQD/exo were treated to each well with a concentration of 0.2 × 10^3^ exosomes per cell. Then, cells were incubated for 6 h at 37°C under 5% CO_2_. To observe the lysosomal uptake of exosomes, *in vitro* experiments were conducted as follows: Cells were cultured at 5 × 10^4^ cells per well in 8-well confocal plate and incubated overnight at 37°C under 5% CO_2_. Three different types of *D-* GQD/exo were treated to each well with a concentration of 0.2 × 10^3^ exosomes per cell. Then, cells were incubated for 4 h at 37°C under 5% CO_2_ and LysoView 594 was added for staining lysosomes in the cells. To observe the direct membrane fusion of exosomes, *in vitro* experiments were conducted as follows: Cells were cultured at 5×10^4^ cells per well in 8-well confocal plate and incubated overnight at 37°C under 5% CO_2_. Hoechst 33342 were added for staining nuclei of the cells. Three different types of *D*-GQD/exo stained with DiI were treated to each well with a concentration of 0.4 × 10^3^ exosomes per cell, followed by incubation for 1 h at 37°C under 5% CO_2_. Then, cellular membranes were stained by DiO. To observe the exosomal cargo (*D*-GQDs) delivery of exosomes, *in vitro* experiments were conducted as follows: Cells were cultured at 5×10^4^ cells per well in 8-well confocal plate and incubated overnight at 37°C under 5% CO_2_. Three different types of *D-*GQD/exo stained with PKH26 were treated to each well with a concentration of 0.2 × 10^3^ exosomes per cell. With the incubation for 6 h at 37°C under 5% CO_2_, cellular membranes were stained by CellBrite Green Cytoplasmic Membrane dye. The SigmaPlot 10.0 was used to generate plots of the analyzed data.

### 2.8. Image Processing and Quantification

All CLSM imaging processing and analytical quantification used the ImageJ software. Prior to every analysis, the manual conversion of length units from pixels to μm by the scale bar on CLSM images should be set. The cellular uptake of exosomes was quantified as follows: Z-stack images of each group were scanned into three z-stack-set with a step size of 0.5 μm. The cellular taken up *D-*GQDs, exosomal cargo markers, were quantified with the integrated intensity analysis. The RawIntDen (RID; a.u.) value which is the sum of all fluorescent intensity in the region-of-interest (ROI) was measured from the montages consisting of CLSM three z-projected images in the blue channel. The red area of cellular cytosol was converted to monochrome (black and white) and the middle (second) image of z-projection was selected for quantification. The cell cytosol area was quantified with the following equation:

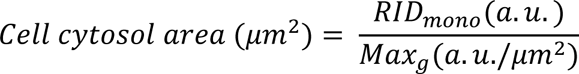

The maximum and minimum gray levels (Max_g_, Min_g_; a.u./ μm^2^) reflect maximum-gray (represented as black) and minimum-gray (represented as white), respectively, and the Min_g_ is zero. Finally, the fluorescence intensity per cell cytosol area was calculated with the following equation:

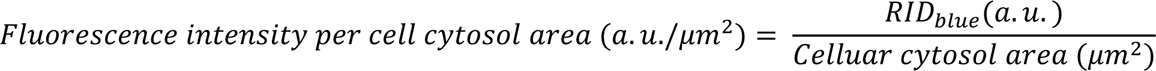

The lysosomal entrapment of exosomes was quantified as follows: Only colocalized purple spots were selected, whereas all non-colocalized red and blue spots were deselected, via ImageJ software with JACOPx Plugin. Then, the purple-colocalized spots were counted. The total counted numbers were divided by the total cell number within scanned images.

The membrane fusion of exosomes was quantified as follows: Pearson’s correlation coefficient (PCC) was calculated for the analytical quantification of colocalization between cellular and exosomal membranes. The scatter plot of the red and green channels can be generated using ImageJ software with the JACOPx Plugin. Subsequently, the PC linear plot can be created based on the scatter plots of each channel. Finally, the PCC was calculated with the following equation:

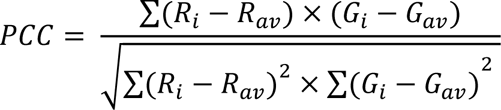

In the entire image, R_i_ and R_av_ represent the intensity values and the average intensity of the red channel, respectively, while G_i_ and G_av_ represent the intensity values and the average intensity of the green channel, respectively. PCC ranges from −1 to 1: −1 for a perfect negative correlation (inverse colocalization); 0 for no correlation (no colocalization); 1 for a perfect positive correlation (full colocalization).^39^ A higher positive PCC value indicates more colocalization. The SigmaPlot 10.0 was used to generate plots of the analyzed data.

### 2.9. Exosomal Protemoics

Western blot was carried out as follows: Collected exosomes were lysed by adding 1X RIPA buffer, and 20 μg of proteins from whole exosome lysates were separated by Sodium dodecyl-sulfate polyacrylamide gel electrophoresis (SDS-PAGE). The separated proteins were transferred to a nitrocellulose membrane (1662807; Bio-Rad, CA, USA) and treated with primary antibodies (anti-CD9 antibody; anti-CD63 antibody; anti-CD81 antibody; anti-CD9 antibody; anti-neuregulin1 antibody; anti-GalNAc antibody; anti-TGF-beta1 antibody; anti-beta-actin antibody). Secondary antibodies (anti-mouse HRP-linked antibody; anti-rabbit HRP-linked antibody) were then blotted and immunoreactive species were detected by an ECL substrate using a bioimaging system. Relative expression levels of detected proteins were quantified using ImageJ software. Sample preparation for MS-based proteomics was as follows: Cell-derived exosomes were lysed by 1X RIPA buffer. 50 ug of protein from exosomes were trypsinized and eluted. Eluted peptides were dried down and resuspended, followed by loading in the ultra-high-resolution MS coupled to nano-ultra-high-pressure LC or CE for bottom-up proteomics. The GraphPad Prism 8.0 (GraphPad Software Inc, SD, USA) and SigmaPlot 10.0 were used to generate plots of the analyzed data.

### 2.10. FRET and its Inhibition in Membrane Fusion

Cells were cultured at 5 × 10^4^ cells per well of 8-well confocal plate and incubated overnight at 37°C under 5% CO_2_. *D*-GQD/exo stained with DiI were treated to each well with a concentration of 0.4 × 10^3^ exosomes per cell, followed by incubation for 1 h at 37°C under 5% CO_2_. Then, cellular membranes were stained by DiO. Cells were washed, trypsinized, and then collected on a 96-well black plate. Fluorescence intensity was measured before and after cell lysis with Tween-20 using a plate reader. The cell suspension was excited at 484 nm, the DiO excitation wavelength, and the fluorescent emission was measured at 565 nm, the DiI emission wavelength. The SigmaPlot 10.0 was used to generate plots of the analyzed data.

## 3. RESULTS AND DISCUSSION

### 3.1. Exosomes from Various Cell-of-Origin

To investigate the distinct cellular uptake and subsequent cargo release profiles of exosomes from various cell-of-origin, three cell lines were selected for this study: HepG2 cells (human hepatoma), 3T3 cells (mouse fibroblast), and HeLa cells (human cervical adenocarcinoma). Exosomes from culture media of these cell lines were isolated using size-based ultrafiltration techniques. We observed the size and the morphology of exosomes using TEM, which confirmed that all isolated exosomes displayed a spherical shape with structural integrity (Figure 1a). Moreover, based on the quantitative analysis of TEM images, the mean diameters of all three types of exosomes were around 100 nm: 101.3 ± 13.2 nm for HepG2 cell-derived exosomes (HepG2 exo), 99.2 ± 16.8 nm for 3T3 cell-derived exosomes (3T3 exo), and 103.3 ± 15.5 nm for HeLa cell-derived exosomes (HeLa exo). To estimate the concentration and size distribution of exosomes, NTA was conducted on exosome suspension samples (Figure 1b). The concentrations of exosomes isolated from 600 mL of serum-free media resulted in 2.80 ± 0.41 × 10^10^ particles/mL for HepG2 exo, 2.62 ± 0.12 × 10^10^ particles/mL for 3T3 exo, and 2.98 ± 0.35 × 10^10^ particles/mL for HeLa exo. We observed a narrow size distribution, and the peaks of diameter were 129.7 ± 4.5 nm for HepG2 exo, 130.1 ± 0.4 nm for 3T3 exo, and 136.7 ± 9.7 nm for HeLa exo. The diameters of exosome measured by NTA appeared to be slightly larger than that of TEM image analysis. This discrepancy was attributed to the specific measurement techniques that were operated in different ways. Using TEM, individual exosome particles were imaged under dehydrated and vacuum conditions, resulting in the observation of shrinkage in exosome size.^40^ Moreover, the negative staining conventionally used for TEM imaging samples has been reported to potentially induce distortion and flattening of particle structures, leading to an underestimation of their actual size.^41^ On the other hands, the exosomes were measured in solution by NTA that track the individual particles based on their Brownian motion.^40^ In NTA, the smaller exosomes might not be detectable due to the limitations of light scattering which led to a shift in the estimated size distribution to a slightly larger range.^42^ Therefore, when characterizing the size of exosomes, the disparity in size distribution needs to be taken into consideration and confirm to ensure that the respective measurements overlap. To evaluate the colloidal stability, we measured the zeta potential of exosomes from the three origins. This is based on the fact that particles with a higher zeta potential display improved stability due to enhanced electrostatic repulsion, effectively hindering aggregation or coagulation.^43^ The zeta potentials were measured by Zetasizer: −9.2 ± 1.0 mV for HepG2 exo, −11.5 ± 1.8 mV for 3T3 exo, and −18.8 ± 0.3 mV for HeLa exo (Figure 1c), proportional to those of their parent cells (Figure S1e). Analyzing the NTA data along with zeta potentials, we observed a correlation between the negative charge of exosomes and the characteristics of their size distribution. HeLa exo were found to have the most negative surface charges among the three types of exosomes, and they exhibited a size distribution with a sharp and high peak, suggesting that a significant portion of the exosomes shared a similar size. On the other hand, HepG2 exo had a less negative surface charge compared to the other exosomes, and they exhibited a lower and wider distribution of sizes, as indicated by a less pronounced peak in the size distribution. This is attributed to the surface charge that contributes to repulsion between particles, thereby leading to a more dispersed and organized arrangement, resulting in a higher concentration of exosomes around a specific size.^44^ Overall, the integrity and colloidal stability of all three types of exosomes were confirmed. Additionally, to further verify that the isolated nanoparticles were indeed exosomes, we confirmed the expression of exosomal biomarkers by conducting a Western blot analysis of CD9, CD63, and CD81, which are commonly enriched in the exosome membrane.^45^ The Western blot analysis of exosome lysates detected clear immunoblotted bands for CD9, CD63, and CD81 for all isolated and purified exosomes from three cell lines (Figure 1d).

**Figure 1.**
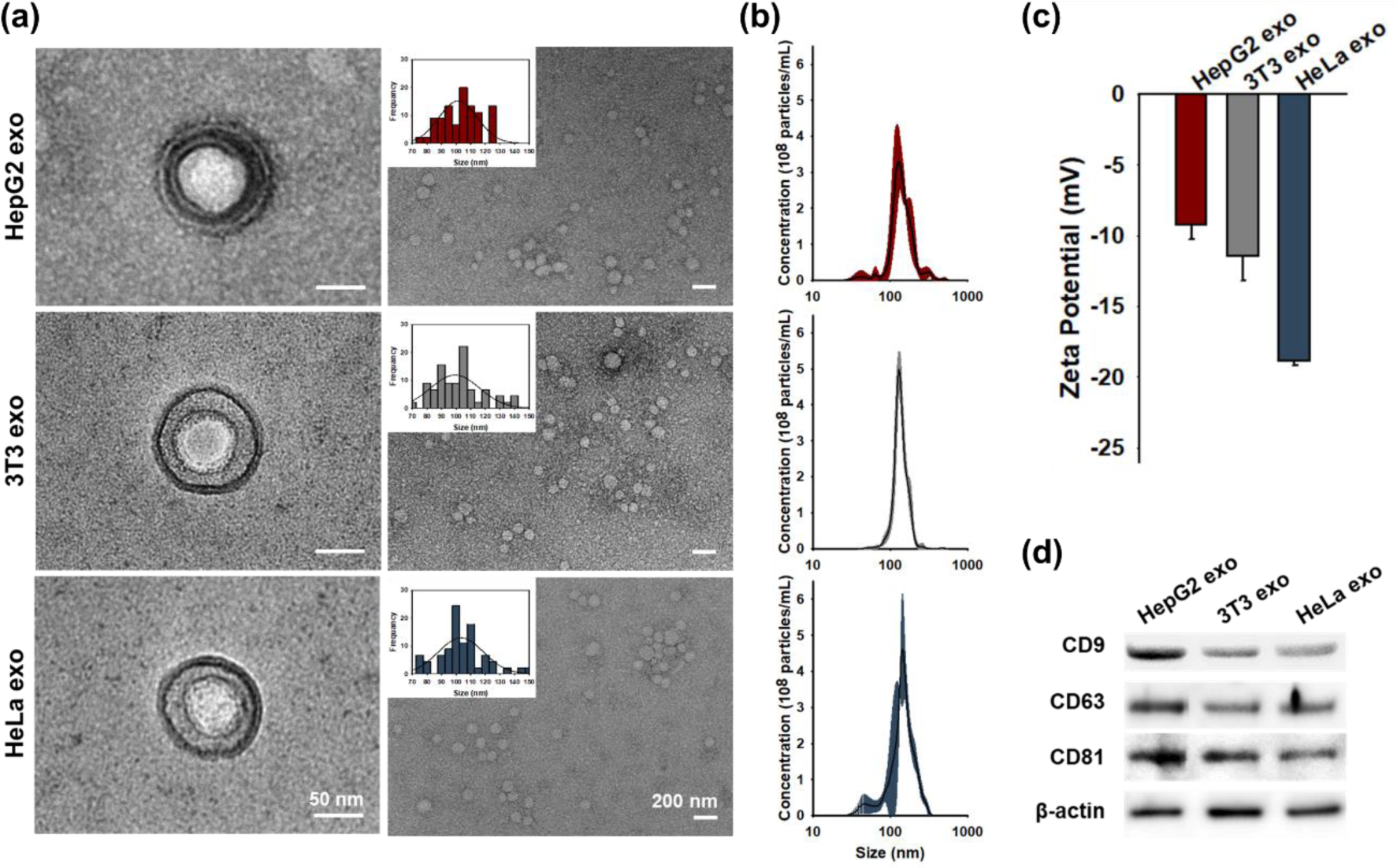
Characterization of exosomes from three different cell lines: HepG2, 3T3, and HeLa cells. (a) Transmission electron microscope (TEM) images of exosomes and size distribution of exosomes based on TEM images. (b) Size distribution and particle number of exosomes measured by a nanoparticle tracking analysis (NTA). (c) Surface potential of exosomes analyzed by a Zetasizer. (d) Western blot analysis of exosomal markers expression.

### 3.2. *D*-GQDs as Representative Cargos in Exosomes

In this study, we utilized chiral GQDs, specifically D-cysteine functionalized GQDs (*D*-GQDs) as representatives cargo in exosomes (Figure 2a and S2). The *D*-GQDs displayed high permeability into exosomes without disrupting the membrane integrity of exosomes due to chiral interaction between *D*-GQDs and biological lipid membrane, and have been developed for exogenous drug loading into exosomes in our previous work.^34, 35^ To confirm the permeation of *D*-GQDs into exosomes, we labeled the exosomes with a red fluorescent membrane dye (PKH26). Meanwhile, *D*-GQDs exhibited strong blue fluorescence emission with the maximum emission wavelength at 450-520 nm when excited at 360 nm (Figure S2e). These optical properties of *D*-GQDs allowed us to observe the permeation of *D*-GQDs into exosomes using CLSM at an excitation wavelength of 405 nm and an emission filter at wavelength of 450 nm. Thus, the colocalization of PHK26-labeled exosomes (red) and *D*-GQDs (blue) indicated the permeation and accumulation of *D*-GQDs inside exosomes (Figure 2b). To ensure the sufficient permeation of *D*-GQDs as representative cargos into exosomes, we quantified the permeation efficiency using the previously developed counting method (See Method).^35^ We quantified that *D*-GQDs (12 μM) can permeate into all three types of exosomes (at a concentration of 1 × 10^9^ particles/mL) with high permeability: 67.2 ± 9.7% for HepG2 exo, 70.6 ± 10.8% for 3T3 exo, and 66.7 ± 4.3% for HeLa exo (Figure 2c and 2d), and achieved 80% by loading them with the optimal concentration of 15 μM *D*-GQDs into exosomes (Figure S3). These efficiencies are higher than any other available drug encapsulation efficiency,^46, 47^ and thus sufficient for *D*-GQDs as representative cargos in the following studies. To note, specific points in CLSM images ranging from 100 to 500 nm appeared larger than the actual size of the objects they represent, which are in the range of 30 to 200 nm. This discrepancy can be attributed to diffraction effects and the inherent resolution limitations of the optical microscope, 180 nm laterally and 500 nm axially,^35, 48^ which are consistent with our previous report.^35^ The permeation of *D-*GQDs into exosomes was further quantified using fluorometry. Fluorescence recovery (Figure S4) became evident in all three types of exosomes loaded with *D*-GQDs after lysis with the biological surfactant, Tween-20. This observation indicated that the self-quenching of *D*-GQDs^49^ within the exosomes was alleviated through the lysis of the exosomal membrane, enabling a sufficient distance for them to enhance their fluorescence without being subjected to the quenching effects of neighboring particles in the lysed suspension.

**Figure 2.**
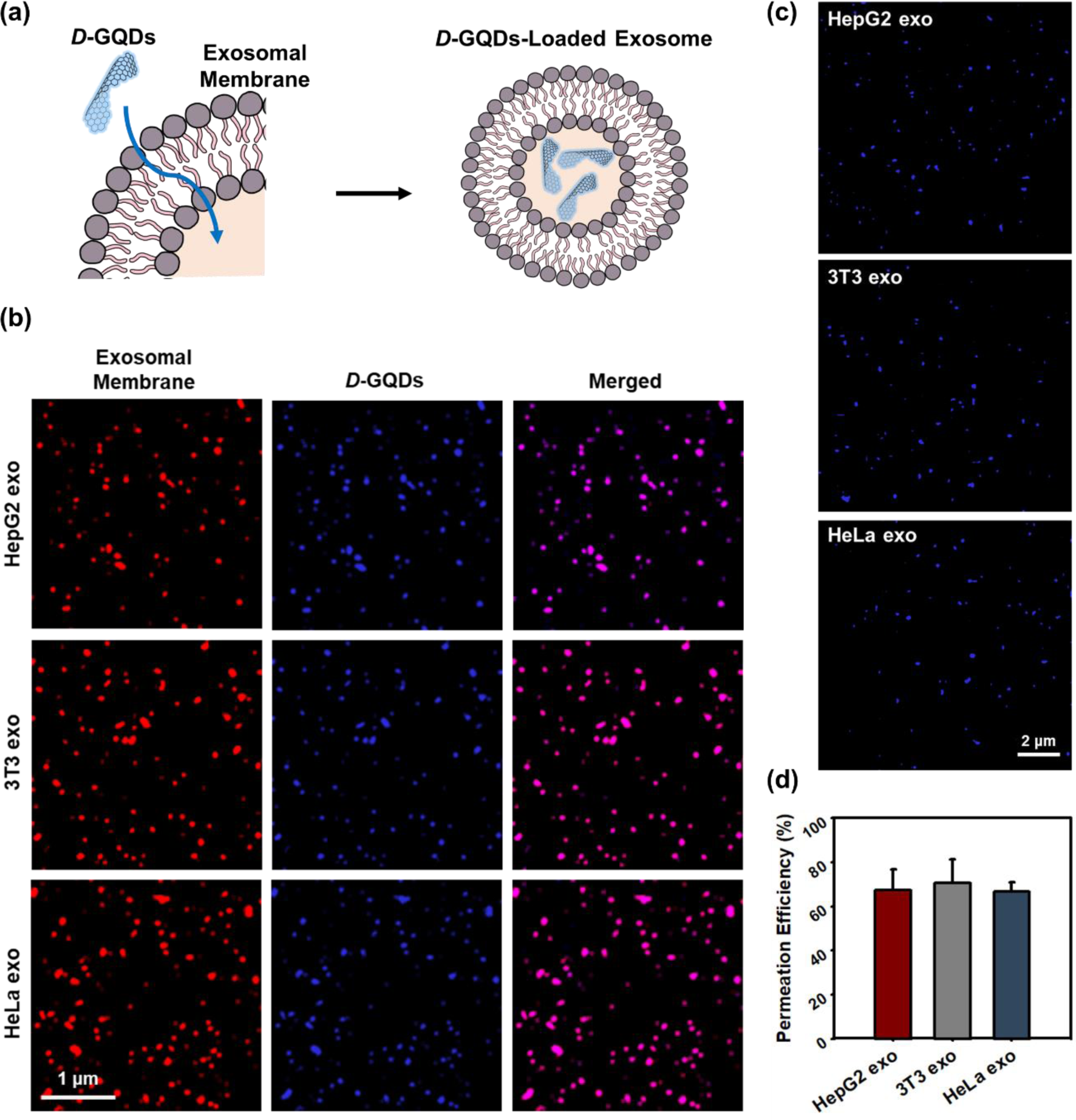
*D*-GQDs as representative of cargos of exosomes. (a) Schematic illustration of the *D*-GQDs permeation into exosomes. (b) Confocal Laser Scanning Microscopy (CLSM) images of *D*-GQDs (blue) loaded PKH26-labeled exosomes (red). (c) Permeation of *D*-GQDs (blue) into exosomes was observed by CLSM. (d) Permeation efficiency was quantified by counting *D*-GQD-loaded exosomes over the total number of exosomes. The samples were prepared by loading 12 μM *D*-GQDs with exosomes (1 × 10^9^ particles/mL), followed by washing with 1X PBS under the support of 100 kDa centrifugal filter tubes.

### 3.3. Differential Cellular Uptake of Exosomes Depending on Cell-of-Origin

Exosomes stemming from various cell types exhibit significant heterogeneity, profoundly influencing their uptake behavior within recipient cells.^50^ To explore the cellular uptake tendencies of exosomes depending on their cell-of-origin and recipient cells, we conducted quantitative analysis of CLSM images. The exosomes were loaded with *D*-GQDs, enabling the tracking through blue fluorescence, while the recipient cells were stained with cytol-tracible red dye. After 6 hours of co-incubation, the cellular uptake of exosomes into their parental or non-parental cells was imaged using CLSM (Figure 3). In order to quantitatively analyze the cellular uptake tendencies of exosomes, we meticulously examined the distribution of *D*-GQDs within the recipient cells by using the equation we developed (See Method). In the recipient group of HepG2 cells, we verified that the uptake of HepG2 exo was significantly higher compared to the other two types of exosomes, being ∼3.2 times higher than 3T3 exo and ∼2.4 times higher than HeLa exo (Figure 3a). Similarly, in the recipient group of 3T3 cells, we observed the highest uptake of 3T3 exo, which was ∼2.2 times higher than and ∼1.4 times higher than HepG2 exo and HeLa exo, respectively (Figure 3b). The consistent trend was also observed in the group of HeLa cells, with uptake of HeLa exo being approximately 2 times higher than HepG2 exo (∼1.7 times) and 3T3 exo (∼1.9 times) (Figure 3c). These results not only agreed with prior findings signifying the propensity of exosomes to home in on their cell-of-origin^17^ but also provided a quantitative comparison of their cellular uptake efficiency in both parental and non-parental recipient cells. In addition to the thorough observation of exosome uptake trends and its quantitative analysis, getting a comprehensive understanding of the intricate mechanisms governing their entry into recipient cells is necessary to to reveal the underlying cellular uptake and subsequent mechanisms.^12^ In previous studies, the cellular uptake of exosomes has been reported to mainly occur through two primary pathways: endocytic uptake and direct membrane fusion.^18, 19, 51–54^ Different pathway will also lead to distinct cargo release mechanisms: lysosomal entrapment followed by degradation^55^ and direct cargo entry into the cytosol.^56^ To further investigate the understanding of distinct uptake efficiency and subsequent cargo release, we studied dissecting the potential uptake pathways of exosomes, as well as unraveling the processes involved in the delivery of their cargo through tracking *D*-GQDS in subsequent experiments.

**Figure 3.**
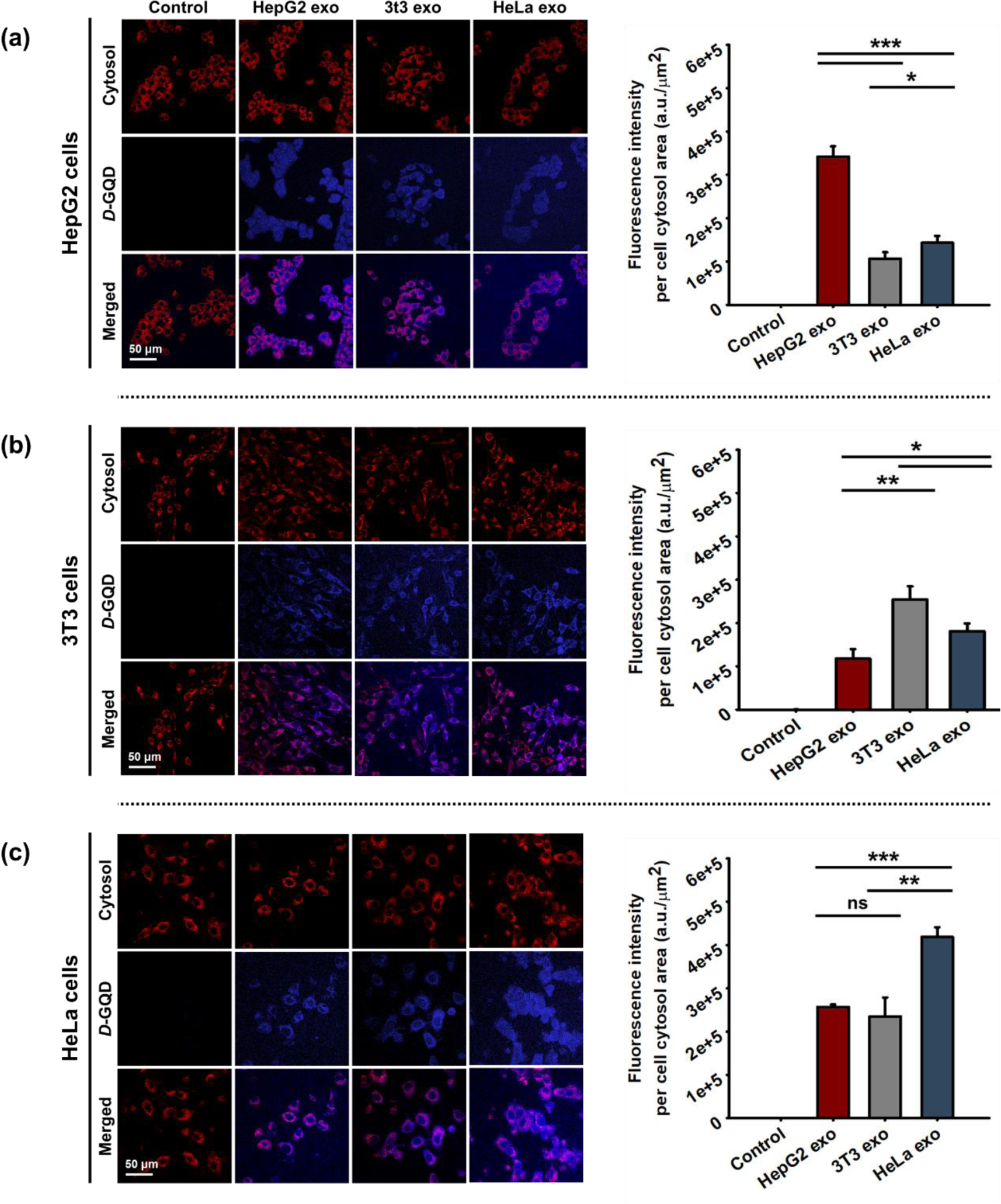
Cellular uptake profiles of *D*-GQDs loaded exosomes derived from three different cell lines into (a) HepG2 cells, (b) 3T3 cells, and (c) HeLa cells (mean ± s.d.). ns: not significant, *: p < 0.05, **: p < 0.01, ***: p < 0.001, ****: p < 0.0001. Each channel represents: red for the cell cytosol area, and blue for *D*-GQDs. HepG2, 3T3, and HeLa cell lines were incubated for 6 hours with cell culture medium (Control) with a concentration of 0.2×10^3^ exo/cell. The samples were prepared by loading 12 μM *D*-GQDs with exosomes (1 × 10^9^ particles/mL).

### 3.4. Endocytosis via Cell Receptors-Exosome Ligands Interaction

To investigate the endocytic pathways involved in the uptake of exosomes by their parental/non-parental cells, as well as the releasing sites of their cargo, we conducted in-depth cellular imaging using CLSM. We used LysoView dye to stain the lysosomes of the recipient cells. This enabled us to observe how exosomes, upon entering the endocytic pathway,^57^ become trapped in endosomes, eventually leading to their cargos entrapped in lysosomes. We incubated exosomes and cells for 4 hours to ensure a sufficient uptake of exosomes by the cells through the endocytic process.^58^ We quantified the colocalized spots of lysosomes and *D*-GQDs to analyze the entrapment of exosomes within lysosomes in recipient cells. Remarkably, our results demonstrated that the amount of intraspecies exosomes entrapped within lysosomes were significantly higher than cross-species exosomes (Figure 4a). HepG2 exo was found to be entrapped within lysosomes of HepG2 cells ∼2.1 times more than 3T3 exo and ∼1.9 times more than HeLa exo (Figure 4b). Similarly, in HeLa cells, the number of colocalization spots for HeLa exo was significantly higher, approximately 2 times higher than HepG2 exo (∼1.7 times) and 3T3 exo (∼2 times) (Figure 4d). Although the colocalization in 3T3 cells was relatively low compared to the other two cancer cell lines, HepG2 and HeLa cells, the consistent trend was observed. 3T3 exo still exhibited colocalization with lysosomes more than ∼1.6 times than HepG2 exo and HeLa exo (Figure 4c). These results indicated that while intraspecies exosomes are preferentially taken up into their parental cells, a large number of exosomes were involved in endocytic uptake, leading to their entrapment in lysosomes.

**Figure 4.**
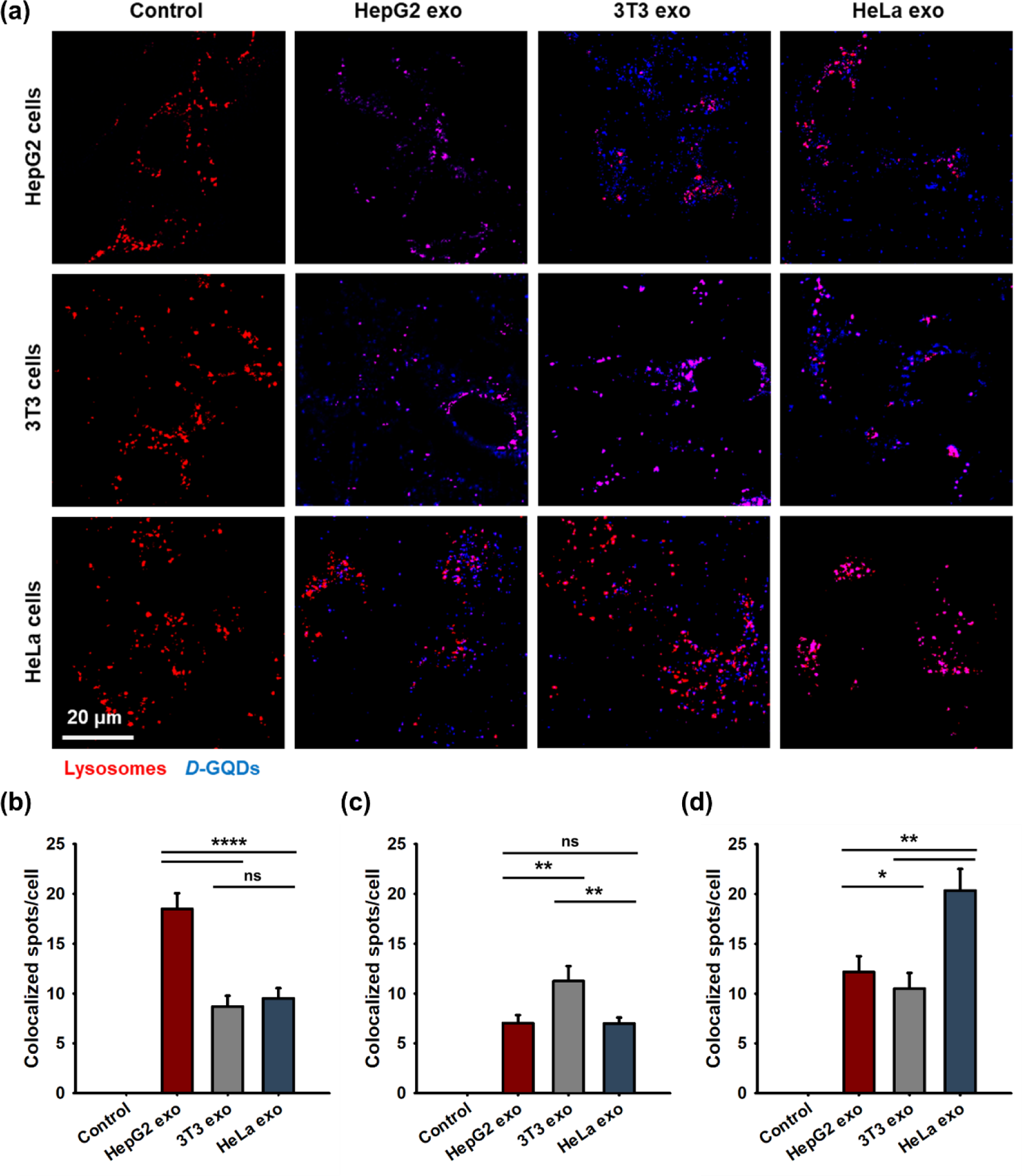
Endocytic uptake profiles of exosomes. (a) CLSM images of three cell lines incubated with exosomes derived from different cell lines. Each channel represents: red for the lysosomes, and blue for *D*-GQDs. The quantification of colocalization between *D*-GQDs and lysosomes was analyzed in (b) HepG2 cells, (c) 3T3 cells, and (d) HeLa cells (mean ± s.e.). ns: not significant, *: p < 0.05, **: p < 0.01, ***: p < 0.001, ****: p < 0.0001. HepG2, 3T3, and HeLa cell lines were incubated for 4 hours with cell culture medium (Control) with a concentration of 0.2 × 10^3^ exo/cell. The samples were prepared by loading 12 μM *D*-GQDs with exosomes (1 × 10^9^ particles/mL).

Endocytosis is primarily mediated by receptor-ligand interactions.^59^ To identify the specific ligands involved in the endocytic uptake pathway with their parental cells, we conducted further investigations. By comparing the analysis of the surface protein composition of exosomes using Mass Spectrometry (MS), we identified and confirmed the presence of distinct proteins on three different exosomes that played a crucial role in mediating receptor-ligand interactions with their parental cells, facilitating the endocytic uptake. Based on the analysis of MS-based proteomics data, we classified several protein accessions into eight distinct proteins: 5 protein accessions for Transforming growth factor-beta 1 (TGF-β1), 13 protein accessions for Glycoproteins (without or with N-acetylgalactosamine (GalNAc) residues, 8 protein accessions and 5 protein accession, respectively), 1 protein accession for Neuregulin 1 (NRG1), 1 protein accession for Beta-actin (β-actin), 3 protein accessions for Clathrin Heavy Chain 1 (CHC1), 4 protein accessions for Heat Shock Protein 70 (HSP70), 6 protein accessions for Heat Shock Protein 90 (HSP90), and 1 protein accession for vinculin. TGF-β1 is a versatile cytokine that regulates various cellular functions, including a wide range of growth factors that control cell growth, differentiation, and other essential cellular processes.^60^ When TGF-β ligands bind to the TGF-β receptor complex, consisting of type I TGF-β receptor (TβRI) and type II TGF-β receptor (TβRII), it triggers a series of endocytic uptake processes.^61^ Glycoproteins are proteins with attached carbohydrate molecules found in living systems, including cell membranes and they play vital roles in cellular functions.^62^ According to the previous reports, glycoproteins with terminal galactose (Gal) and/or N-acetylgalactosamine (GalNAc) residues exhibit high specificity and affinity for the liver-specific Asialoglycoprotein receptor (ASGPR), leading to their internalization through endocytosis.^63, 64^ Specifically, in HepG2 cells, a significant overexpression of ASGPR on the cell membranes was demonstrated, whereas notably lower ASGPR expression was observed on HeLa cells.^65^ NRG1 is a signaling protein that regulates cell-to-cell communication and plays critical roles in cell function.^66^ NRG binds to the Epidermal Growth Factor Receptor (EGFR), which includes family members ErbB1, ErbB2, ErbB3, and ErbB4, followed by subsequent cellular processes such as endocytosis.^67, 68^ In particular, it was reported that NRG1 directly bind to ErbB3 and ErbB4, and not bind directly but related functionally to ErbB2.^69^ EGFR is reported to be overexpressed in around 90% of cervical tumors, including HeLa cells.^70, 71^ Furthermore, ErbB4 was commonly observed on the surface of HeLa cells.^72, 73^ The interaction between specific ligands (TGF-β1, GalNAc, and NRG1) with their receptors (TβR, ASGPR, and ErbB4) were demonstrated respectively to be followed by subsequent cellular processes, such as clathrin-dependent endocytosis.^64, 67, 74^ These receptor-mediated endocytosis involves clathrin-coated pits and vesicles assembled and formed by CHC.^75^ The detection of CHC1 in all three exosomes provided additional evidence supporting the internalization of specific molecules from the extracellular space through endocytosis. Both HSP70 and HSP90 have been commonly found associated with exosomes, identifying them as exosomal markers.^76^ Beta-actin (β-actin) and Vinculin exhibit stable expression in most cell types and are commonly used as loading controls to normalize protein expression across samples.^77, 78^ The presence of these proteins detected in all exosome samples confirmed the reliable measurement of our MS-based proteomics data.

To confirm the expression levels of TGF-β1, GalNAc, and NRG1, which are closely related to the interaction with their parental recipient cells, we, therefore, performed Western blot analysis (Figure 5b). In order to ensure the accuracy and reliability of quantitative analysis, β-actin was used as the loading control, and the relative expression levels were normalized using β-actin. We observed that the expression level of TGF-β1 on HeLa exo was two-fold higher than that of HepG2 exo at a blotting concentration of 0.3 μg/mL of anti-TGF-β1 antibodies, while no band was observed in 3T3 exo. (Figure 5c). According to the report, both HepG2 and HeLa cells exhibited a high expression level of TβRII, and significant responsiveness to TGF-β1.^79^ Hence, when HepG2 exo and HeLa exo are exposed to cell surfaces expressing TβR, they are likely to undergo the receptor-ligand interaction-mediated endocytic process. Notably, intraspecies exosome groups were observed to be prominently entrapped within lysosomes in both HepG2 and HeLa cells (Figure 4b and 4d). In contrast, no significant entrapment was observed in the intraspecies group in 3T3 cells (Figure 4c). These results were attributed to the interaction between TGF-β1 expressed on exosomes and TβR present in HepG2 cells and HeLa cells. Moreover, we observed the expression of glycoproteins with GalNAc residues on HepG2 exo, while no expression in 3T3 exo and HeLa exo at a blotting concentration of 0.4 μg/mL anti-GalNAc antibodies (Figure 5d). This implies that significant endocytosis will take place through the interaction between GalNAc on HepG2 exo and the overexpressed ASGPR on HepG2 cells. Indeed, the predominant lysosomal uptake of HepG2 exo by HepG2 cells was observed (Figure 4b). Furthermore, consistent with the MS-based proteomics, we detected the expression of NRG1 only in HeLa exo at a blotting concentration of 0.4 μg/mL of anti-TGF-β1 antibodies, while no expression was observed in 3T3 exo and HepG2 exo (Figure 5e). Based on the comprehensive analysis of lysosomal uptake results in HeLa cells, we confirmed that the interaction between NRG1 on HeLa exo and ErbB4 on HeLa cells led to a significant increase in the lysosomal entrapment of HeLa exo compared to HepG2 exo and 3T3 exo (Figure 4d). These experiments illustrated that the intraspecies exosomes carry ligands (TGF-β1, GalNAc, and NRG1) capable of interacting with receptors (TβR, ASGPR, and ErbB4) on their parental cells, resulting in endocytic uptake followed by the entrapment of exosomal cargo within lysosomes.

**Figure 5.**
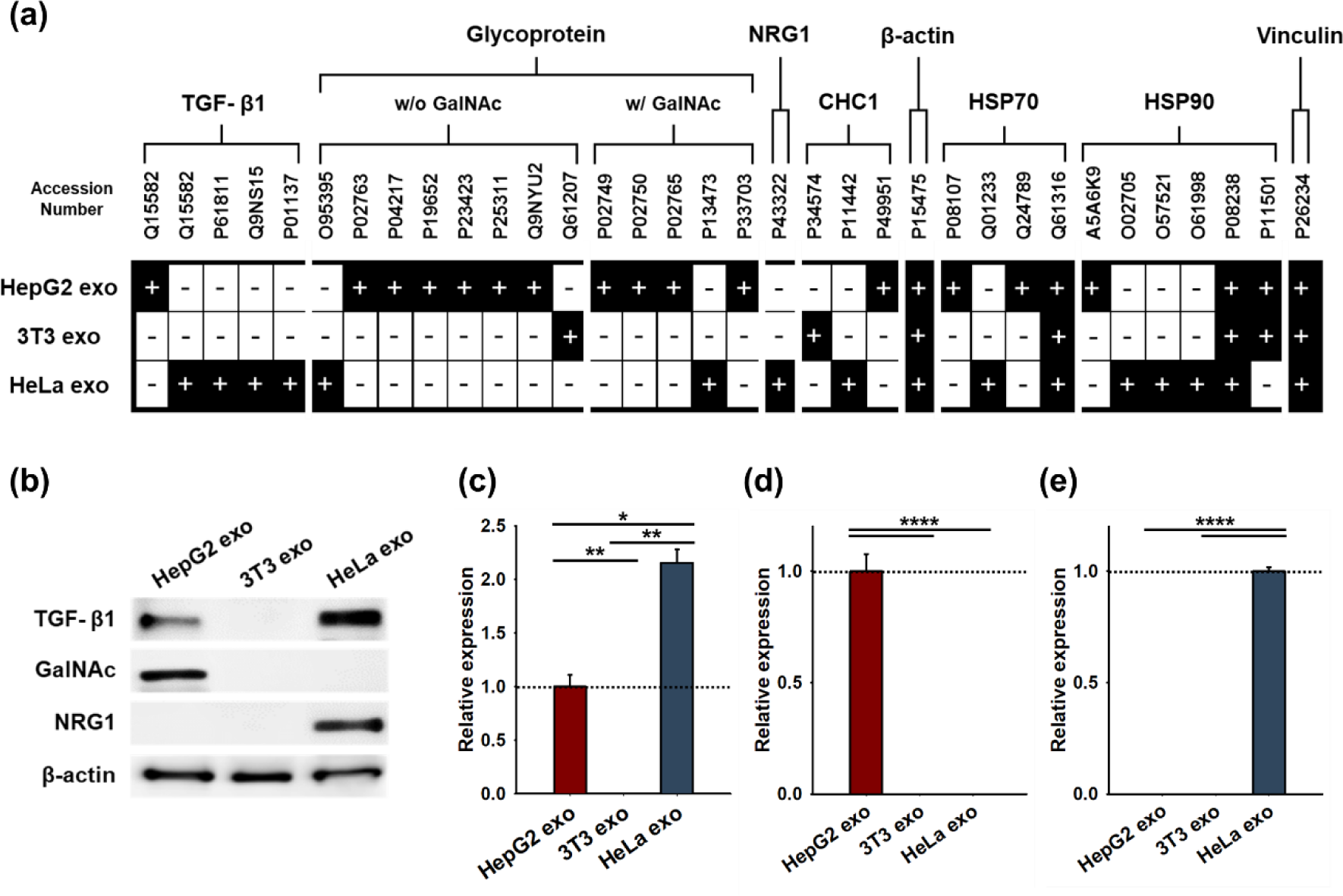
Identification of the surface proteins on exosomes that facilitate the receptor-ligand interaction mediated endocytosis. (a) Mass spectrometer (MS)-based proteomics. Protein accession numbers were retrieved from the UniProtKB/Swiss-Prot. Black boxes with [+] corresponded to the proteins that were detected, while the white boxes with [−] indicates undetected. TGF: Transforming growth factor, GalNAc: N-acetylgalactosamine, NRG: Neuregulin, CHC: Clathrin-Heavy Chain, HSP: Heat shock protein. (b) Western blot analysis of TGF-β1, GalNAc, and NRG1 expression for three types of exosomes. Relative expression levels of (c) TGF-β1, (d) GalNAc, and (e) NRG1 (mean ± s.e.); Data were normalized to the expression level of β -actin. ns: not significant, *: p < 0.05, **: p < 0.01, ***: p < 0.001, ****: p < 0.0001.

### 3.5. Cellular Uptake via Direct Membrane Fusion with Exosomes

Direct membrane fusion is another primary pathway for cellular uptake of exosomes. Hence, to further investigate the underlying mechanism of membrane fusion, we performed colocalization examination and Fluorescence Resonance Energy Transfer (FRET) analysis. First, we introduced a red membrane fluorescent dye, DiI, to label the lipid bilayer of exosomes. After incubation with exosomes for 1 hour, the cells were stained with a green membrane fluorescent dye, DiO. As shown in CLSM images (Figure 6a-c), a significant overlap of cellular membranes (green) and exosomal membranes (red), indicating the exosome fused with cell membrane, was observed in the groups treated with cross-species exosomes. For the quantitative analysis, the obtained CLSM images were compared using Pearson’s correlation coefficient (PCC) which is a statistical measure in colocalization analysis, quantifying pixel intensity similarity in two fluorescence channels to indicate colocalization extent. Higher PCC value indicated more colocalization between the two channel signals: green, representing cellular membranes, and red, representing exosomal membranes. In HepG2 cells, the PCC value for HepG2 exo was slightly higher than that of the negative control that was treated with media only. In contrast, the PCC values for 3T3 exo and HeLa exo were estimated to be over two-fold higher than that of HepG2 exo (Figure 6d). In the same way, in HeLa cells, the control and HeLa exo groups showed only a small correlation value less than 0.4, whereas both cross-species exosomes, HepG2 and 3T3, estimated the strong correlation with the large PCC (>0.7) (Figure 6f). Consistently, we observed a parallel trend in the 3T3 cell group, although the difference in colocalization between intraspecies and cross-species exosomes was less pronounced compared to HepG2 and HeLa cells. Both HepG2 exo and HeLa exo exhibited a strong correlation between the red and green channels, with PCC values greater than 0.7. On the other hand, 3T3 exo showed a moderate correlation value, ranging from 0.4 to 0.7 (Figure 6e). These findings validated that the PCC of cross-species exosomes mostly exhibited large correlation values, indicating a high degree of colocalization between two fluorescent membranes. This observation suggested that the uptake pathway of cross-species exosomes was more likely to involve direct membrane fusion. Additionally, the presence of exosomes (red dots) within intracellular area was clearly visible in the groups treated with intraspecies exosomes across all three cell types. This observation suggested endocytic uptake occurred even after just 1 hour of incubation (Figure 6a-c). These findings provided further evidence of the previously observed trends that intraspecies exosomes were internalized more effectively through the endocytic pathway (Figure 4).

**Figure 6.**
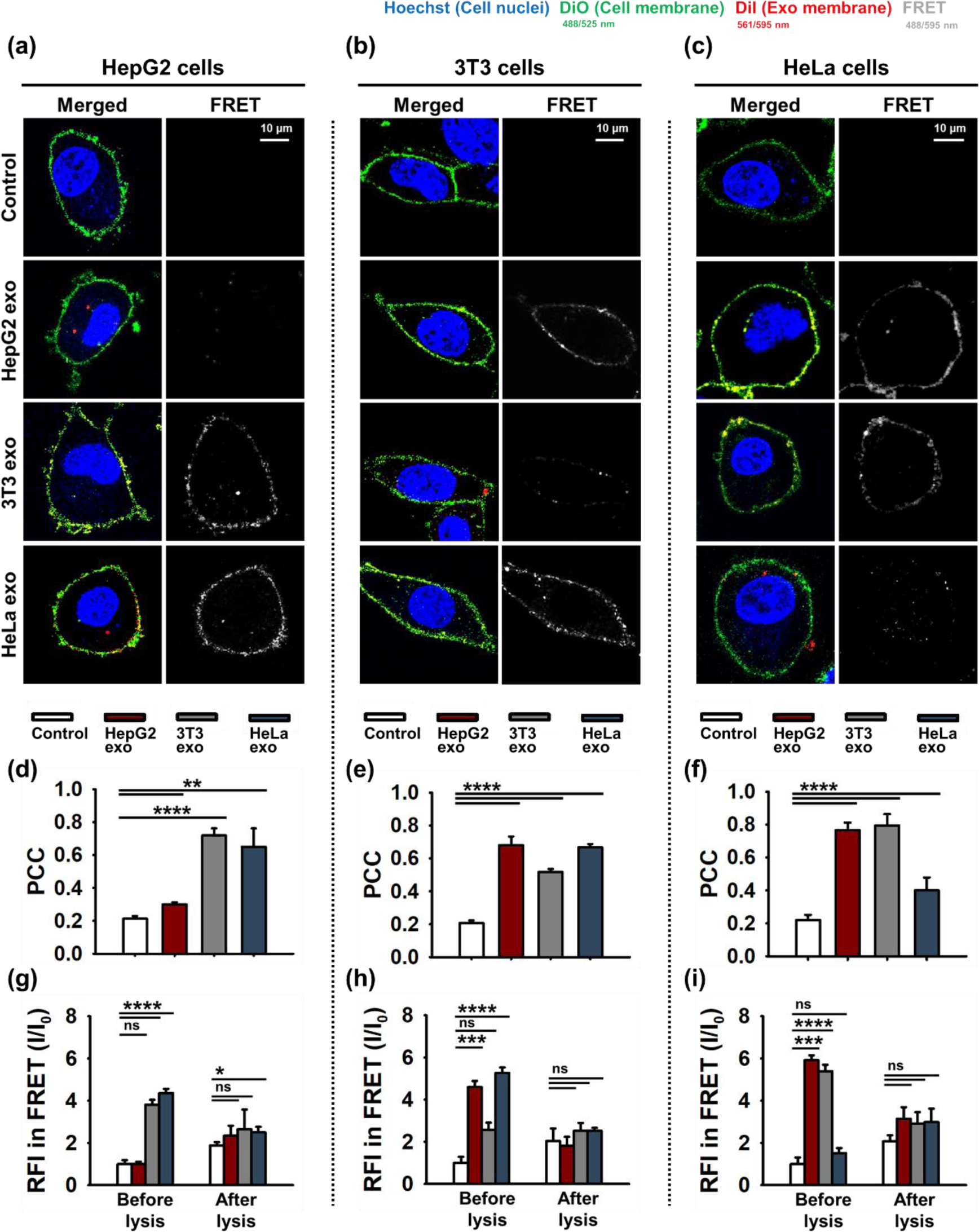
Membrane fusion between cells and exosomes. CLSM images of (a) HepG2 cells, (b) 3T3 cells, and (c) HeLa cells after 1 hour of incubation. Each channel represents: blue for nuclei, green for cellular membrane, red for exosomal membrane, and gray for FRET. (d-f) PCC based on merged images was quantitatively analyzed and (g-i) FRET evaluation by measuring fluorescence intensity (ex: 484 nm, em: 565 nm) was measured in cell suspension (mean ± s.d.). ns: not significant, *: p < 0.05, **: p < 0.01, ***: p < 0.001, ****: p < 0.0001. HepG2, 3T3, and HeLa cell lines were incubated with cell culture medium (Control) with a concentration of 0.4 × 10^3^ exo/cell.

In addition, to confirm that the overlapping of the green and red signals of CLSM images indeed resulted from membrane fusion, we conducted FRET analysis. FRET provides valuable evidence of the proximity and interaction of fluorescent molecules, as it enables us to detect energy transfer between a donor and acceptor fluorophore within a range of 1-10 nm.^80^ In this experiment, DiO was used as the fluorescence donor, and DiI was used as the fluorescent acceptor to measure FRET. In the intraspecies exosome-treated groups, either a minimal or no FRET-mediated fluorescence signal was observed in the CLSM images. On the other hand, in the cross-species exosome treated group, the CLSM images clearly showed significant red fluorescence (represented in gray), which indicated notable membrane fusion (Figure 6a-c). These FRET signals generated from membrane fusion were further validated by measuring fluorescence intensity (FI) using plate reader. After 1 hour incubation, the cell suspension was excited at 484 nm, which corresponded to the DiO excitation wavelength, and the emitted fluorescence was measured at 565 nm, which corresponded to the DiI emission wavelength. The cross-species exosomes exhibited significantly higher FI in FRET, whereas intraspecies exosomes mostly exhibited lower FI in FRET, comparable to the control group or distinctly lower than cross-species exosomes (Figure 6g-i). Afterward, the inhibition of FRET was also carried out by lysing cells and exosomes with Tween-20, and we confirmed the fluorescence intensity change in the “After lysis” groups (Figure 6g-i). The elimination of quenching resulted in an increase in FI for the control and intraspecies exosome group. However, the FRET generated from cross-species membrane fusion was inhibited, leading to a decrease in FI.^81^ We further ensured the effective inhibition of FRET by observing larger error bars in the “After lysis” groups compared to the “Before lysis” groups. These results were coherently supported by the analysis of PCC values. Overall, both CLSM analysis and fluorimetry demonstrated the propensity for membrane fusion in cross-species exosomes.

### 3.6. Different Exosomal Cargo Release Dependent on Uptake Pathways

To trace exosomal cargo, we observed the retention and release of *D*-GQDs in/from exosomes using CLSM when they were taken up by parental/non-parental recipient cells. *D*-GQDs were loaded into exosomes to represent exosomal cargo, and the exosomal membrane was stained with PKH26. After 6 hours of incubation with cells, the cells were stained with a green cytoplasmic membrane dye, enabling the selective imaging of cell membrane boundaries. *D*-GQDs loaded in intraspecies exosomes were seen colocalizing as purple dots within the cells (Figure 7 and S8), implying that exosomes were taken up by parental recipient cells while preserving their structural integrity. This suggested that the cellular uptake of intraspecies exosomes occurred through endocytosis, resulting in the entrapment of exosomes within lysosomes and intercepting the release of cargo. On the other hand, *D*-GQDs loaded in cross-species exosomes were observed escaping from the exosomal membrane, represented by the arrows in Figure 7. Upon the fusion of exosomal membranes with cellular membranes, we observed the immediate release of exosomal cargo without the exosomes being enclosed in other lipid vesicles. This fusion process involves a hemifusion stage, followed by the formation of a fusion pore that facilitates the direct transfer of cargo between exosomes and cells.^82^ From this CLSM images, we observed distinct trends of cargo release that correlate with specific uptake pathways, aligning with the lysosomal uptake trend (Figure 4) and the membrane fusion trends (Figure 6). Comprehensively, we demonstrated that intraspecies exosomes preferentially undergo uptake by their parental recipient cells through endocytosis, leading to cargo entrapment within lysosomes (Figure 8a), whereas cross-species exosomes are more inclined to directly release their cargo through membrane fusions to non-parental recipient cells (Figure 8b).

**Figure 7.**
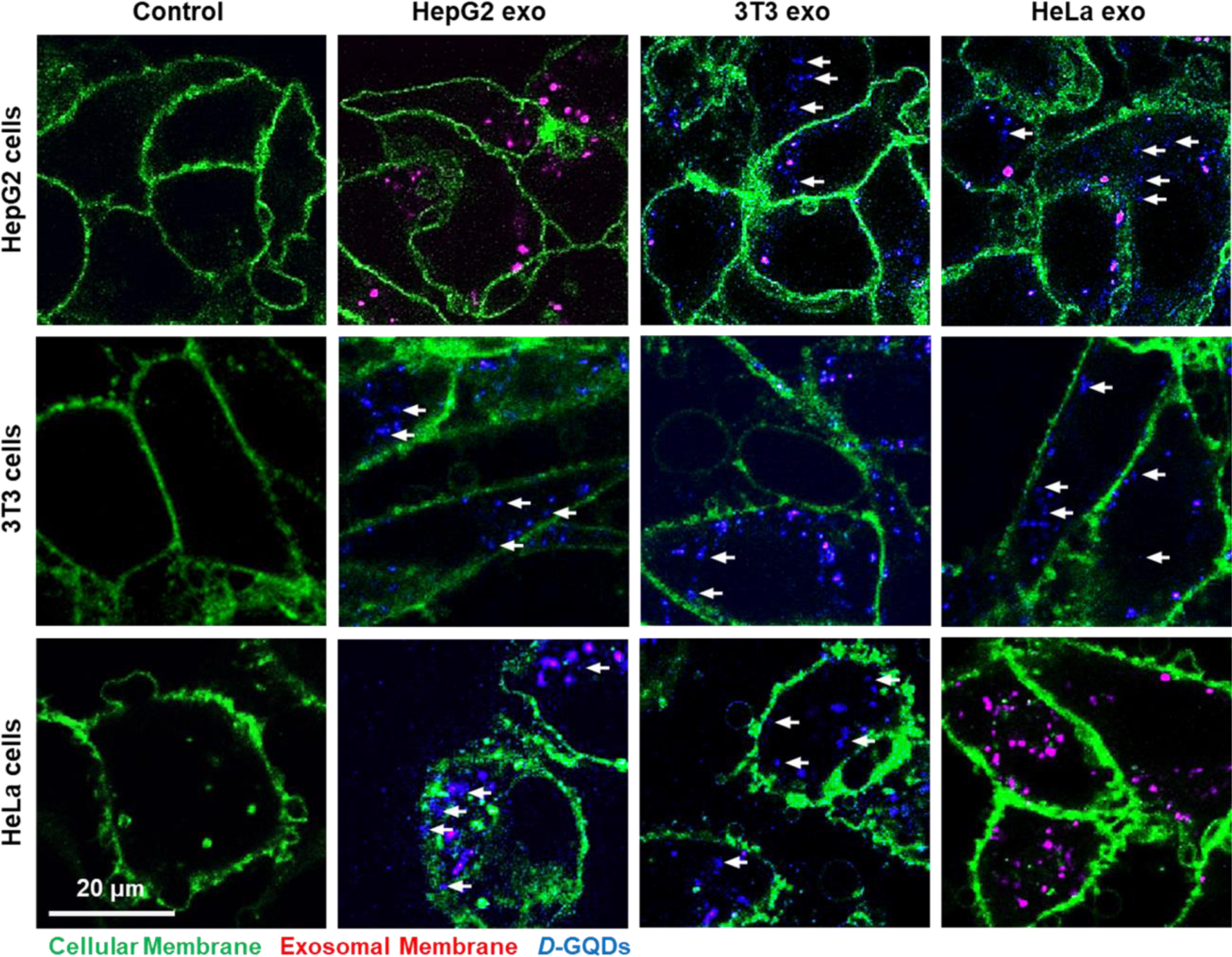
*D*-GQDs release and retention from exosomes after 6 hours of incubation with HepG2 cells (top line), 3T3 cells (middle line), and HeLa cells (bottom line). Each channel represents: green for the cellular membrane, red for the exosomal membrane, and blue for *D-*GQDs. The arrows indicate *D*-GQDs, representing the exosomal cargo released from the exosomes. HepG2, 3T3, and HeLa cell lines were incubated with *D*-GQDs loaded exosomes at a concentration of 0.2 × 10^3^ exo/cell, while the control group was treated with cell culture medium alone. The samples were prepared by loading 12 μM *D*-GQDs with exosomes (1 × 10^9^ particles/mL).

**Figure 8.**
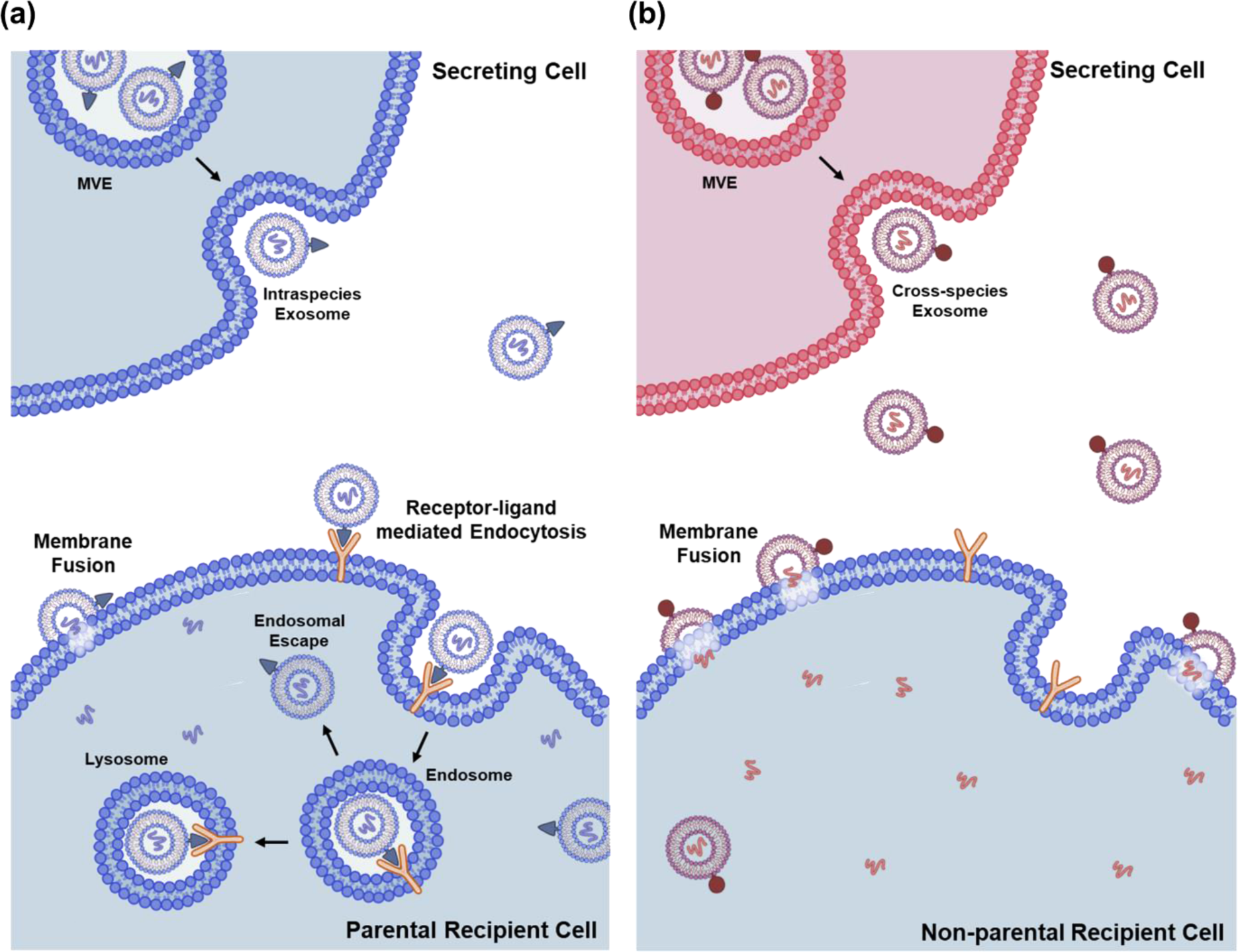
The schematic illustration for the cellular uptake mechanisms of exosomes, depending on their cell-of-origin, followed by cargo release based on the specific cellular uptake mechanisms. (a) Intraspecies endocytic uptake: Exosomes from the same cell-of-origin demonstrated a greater tendency to undergo cellular uptake by parental recipient cells through endocytosis mediated by receptor-ligand interactions, resulting in the entrapment of cargo within lysosomes. (b) Cross-species direct fusion uptake: Exosomes derived from different cells of origin were taken up less by non-parental recipient cells, but primarily through direct membrane fusion, resulting in the direct release of cargo into the cytosol.

### 3.7. Discussion

While the observation of cellular uptake trends of exosomes involving the endocytic pathway and direct membrane fusion has been explored, the detailed mechanisms governing their entry into both parental and non-parental recipient cells are not yet fully understood.^18, 50^ Several approaches have been developed to trace exosomes and their cargo. However, they have some challenges in effectively tracking the journey of exosomes from extracellular interactions to the intracellular release of their cargo. This limitation arises due to inaccuracies resulting from uneven or non-specific staining, coupled with the potential of dyes to modify the cellular membrane properties of exosomes. On top of that, while loading traceable probes into exosomes is of utmost importance to explore intracellular cargo release tracking, the conventional cargo-probe loading approaches face challenges due to their low loading efficiency and the potential risk of damaging the exosomal membrane. Therefore, the lack of comprehension regarding their physiological tendencies, which are involved in their interactions with recipient cells and the intracellular release of their cargo depending on the uptake pathways, leads to a challenge in the effective application of exosomes for drug delivery in clinical uses. In this study, we utilized chiral GQDs to track exosomal cargo, taking advantage of their effective membrane permeability and optical properties while maintaining the structural integrity of the exosomal membrane. By loading *D*-GQDs into exosomes originating from three distinct cell types, we gained a better understanding of mechanisms of exosomal uptake. This process is significantly dependent on both their cell-of-origin and the recipient cell types. Additionally, we conducted further investigations into the subsequent tendency of cargo release based on the specific uptake pathways.

The CLSM results by analyzing the uptake profile of *D*-GQDs-loaded exosomes confirmed that intraspecies exosomes are preferentially taken up by their corresponding parental cells compared to cross-species exosomes (Figure 3 and S5). We further revealed that the enhanced uptake of intraspecies exosomes results from a higher proportion of endocytic uptake facilitated by ligand-receptor interactions (Figure 4 and 5). Notably, in the HeLa cell group, HepG2 exo exhibited a markedly higher lysosomal uptake compared to 3T3 exo, a difference that was statistically significant with a p-value of less than 0.05 (Figure 4d). However, in the HepG2 cell group, HeLa exo did not exhibit a significant difference from 3T3 exo (Figure 4b). These observations implied that HepG2 exo had a greater propensity to interact membrane surface proteins, potentially TβR, on HeLa cells, than HeLa exo does on HepG2 cells. It has been reported that the cleavage of exosome surface proteins resulted in a reduction of their association with recipient cells, confirming the significance of specific proteins that facilitate the cellular uptake of exosomes through an endocytic uptake pathway.^16^

One aspect that requires consideration is that we observed that a large number of intraspecies exosomes were present within lysosomes, rather than their cargo being released into the cytosolic area (Figure 4 and S6). Despite achieving high exosome uptake into their parent cells mainly through receptor-ligand mediated endocytosis, it is necessary to take account the entrapment of exosomes within endosomes, along with their potential subsequent processes of maturation into lysosomes and degradation. Even though endocytosis allows effective cellular uptake of exosomes, this process leads to their subsequent processing and degradation in endosomes and lysosomes, where an acidic environment facilitates various enzymatic activities and is replete with hydrolytic enzymes.^83^ To ameliorate or circumvent undesired degradation and deactivation of therapeutic efficacy within the lysosome, it is necessary to ensure effective drug delivery to intracellular sites without degradation through successful endosome/lysosome escape of exosomes. However, it was reported that only 24 ± 1.2% of intake exosomes were released from endosome/lysosome.^58^ This highlights one of the challenges faced in utilizing pristine exosomes for drug delivery. Therefore, a functionalization approach, such as surface modification on exosomes to facilitate endosomal/lysosomal escape,^84–86^ holds significant potential for advancing therapeutic exosomes. This strategy enables us to fully utilize the inherent homing effect of exosomes for targeted drug delivery.

Another pathway of exosome uptake is direct membrane fusion with the plasma membrane of recipient cells. Cross-species exosomes exhibited a pronounced membrane fusion, whereas the membrane fusion of intraspecies exosomes was less observed (Figure 6 and S7). When we incubated cross-species exosomes with recipient cells, a strong correlation with PCC values larger than 0.7 was observed. In contrast, a weak correlation with PCC values, mostly smaller than 0.4, was seen in the intraspecies exosome-treated groups (Figure 6b-f). Interestingly, the 3T3 cell group exhibited a relatively higher PCC (moderate PCC values of 0.4∼0.7) when incubated with intraspecies exosomes, compared to the HepG2 and HeLa cell groups (Figure 6b, e, and h). Combined with the analysis of overall lower lysosomal uptake counts in 3T3 cells (Figure 4c), these results suggested that despite endocytosis mediated by the receptor-ligand interaction remains dominant for intraspecies uptake of exosomes, non-cancerous cells, such as 3T3 cells, are more likely to be taken up 3T3 exo through membrane fusion compared to cancerous cells such as HepG2 cells and HeLa cells. Direct membrane fusion between lipid membranes is a highly regulated and orchestrated process involving several factors. Especially, the lipid composition of both exosomes and the cell membrane plays a crucial role in fusion. H^+^ associated lipids, such as phosphatidylethanolamine (PE) and phosphatidylserine (PS), were reported to facilitate monovalent cation-induced fusion.^87^ In certain lipid compositions, changes in calcium levels and pH can trigger membrane fusion by destabilizing the lipid bilayers, thereby promoting the fusion process.^88, 89^ The presence of specific proteins known as fusogens, such as SNARE proteins, also influences membrane fusion.^90^ While our study has made progress in understanding cellular uptake of exosomes, the intricate underlying mechanisms that govern the direct membrane fusion of cross-species exosomes still require further investigation. Through the elucidation of factors that influence exosomal membrane fusion, the potential of exosomes for intercellular communication and cargo delivery in various biological contexts can be thoroughly comprehended.

## 4. CONCLUSION

In this study, we utilized fluorescent chiral GQDs to demonstrate not only the preferential uptake pathway of exosomes, depending on their cell-of-origin and recipient cell type, but also the cargo release associated with the specific uptake pathway. We revealed that intraspecies exosomes were mainly taken up by their parental recipient cells through receptor-ligand interaction-mediated endocytosis, leading to their cargo entrapment within lysosomes. On the other hand, despite the lower uptake efficiency, cross-species exosomes were predominantly taken up by non-parental recipient cells through direct membrane fusion, followed by direct cargo release into the cytosol. Our study provides potential scientific significance as we gained better understanding of cellular uptake and intracellular cargo release mechanisms of exosomes. Additionally, we demonstrated membrane fusion, which was previously challenging to elucidate clearly. Our study provides crucial insights that contribute to the progress of effectively utilizing exosomes as drug delivery carriers and enhancing our understanding of intercellular communication.

## AUTHOR INFORMATION

### Corresponding author

*Tel: 574-631-2617. Fax: 574-631-2617.

### Author contributions

YW supervised all the work including overall conceptualization, validation, data analysis, and writing the manuscript. GK designed the methodology and carried out all the experiments. RZ and YZ synthesized and characterized chiral GQDs. GK conducted all CLSM imaging and curated all CLSM data. GK and HJ applied statistical techniques to analyze CLSM data. GK wrote the original manuscript and generated schematic illustrations. All authors contributed to data interpretation, discussion, and writing.

## Acknowledgements

Transmission Electron Microscopy and Confocal Laser Scanning Microscopy images presented in this paper were obtained using the instruments of Notre Dame Integrated Imaging Facility/NDIIF, University of Notre Dame. Mass spectrometer analysis in this paper was measured by the instrument of Mass Spectrometry & Proteomics Facility/MSPF, University of Notre Dame. Western blot images for immunodetection of proteins presented in this paper were acquired using the instrument of Biophysics Instrumentation Core/BIC, University of Notre Dame. We appreciate the support from these facilities for this study.

## Statements and Declarations

### Ethical approval statemen

This study did not involve any experiments conducted on either animals or humans.

