## Supplemental information for "Fluorescent Chiral Quantum Dots to Unveil Origin-Dependent Exosome Uptake and Cargo Release"

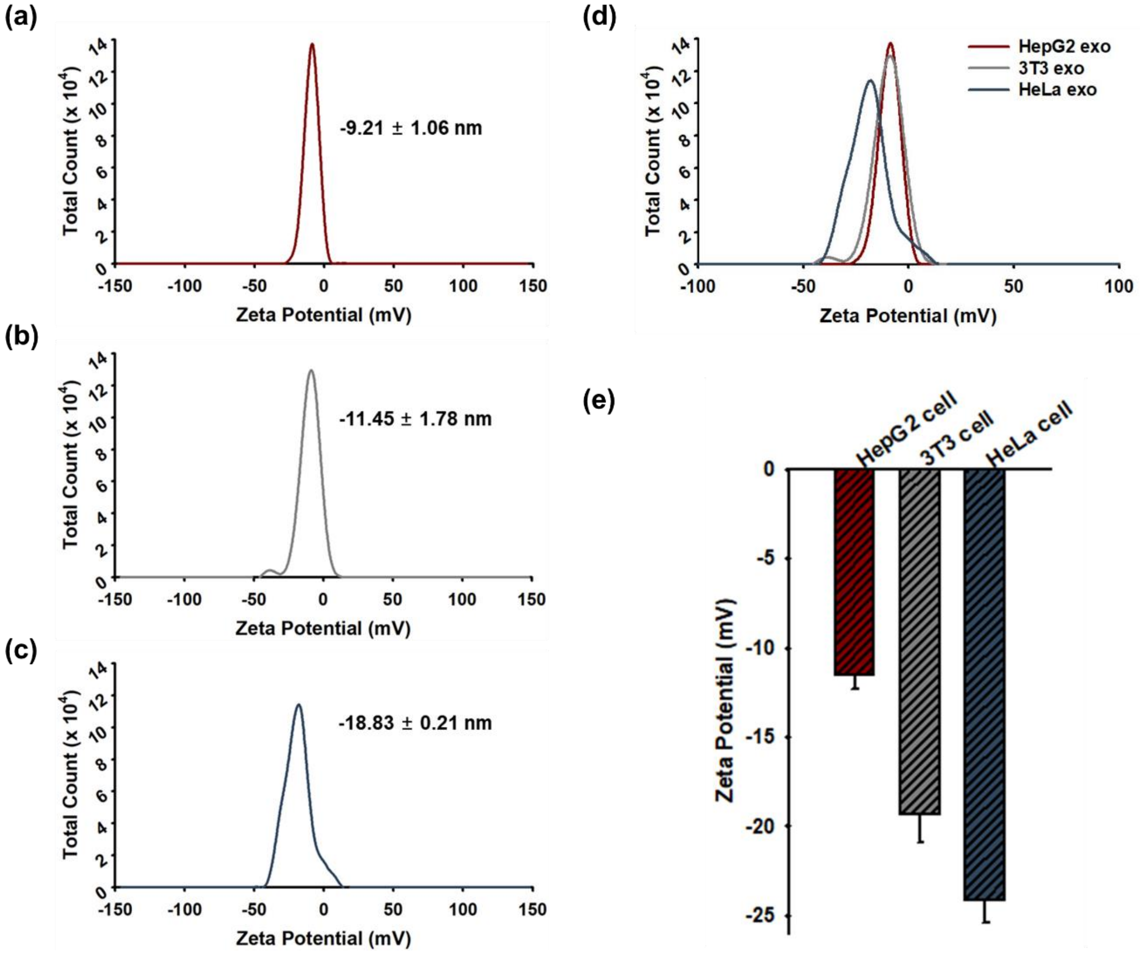


**Figure S1.** Surface potential of exosomes derived from (a) HepG2, (b) 3T3, and (c) HeLa cells were measured individually, followed by plotting (d) the merged result for comparison. (e) The surface charge of their respective cell-of-origin (HepG2, 3T3, and HeLa cells) was also measured.

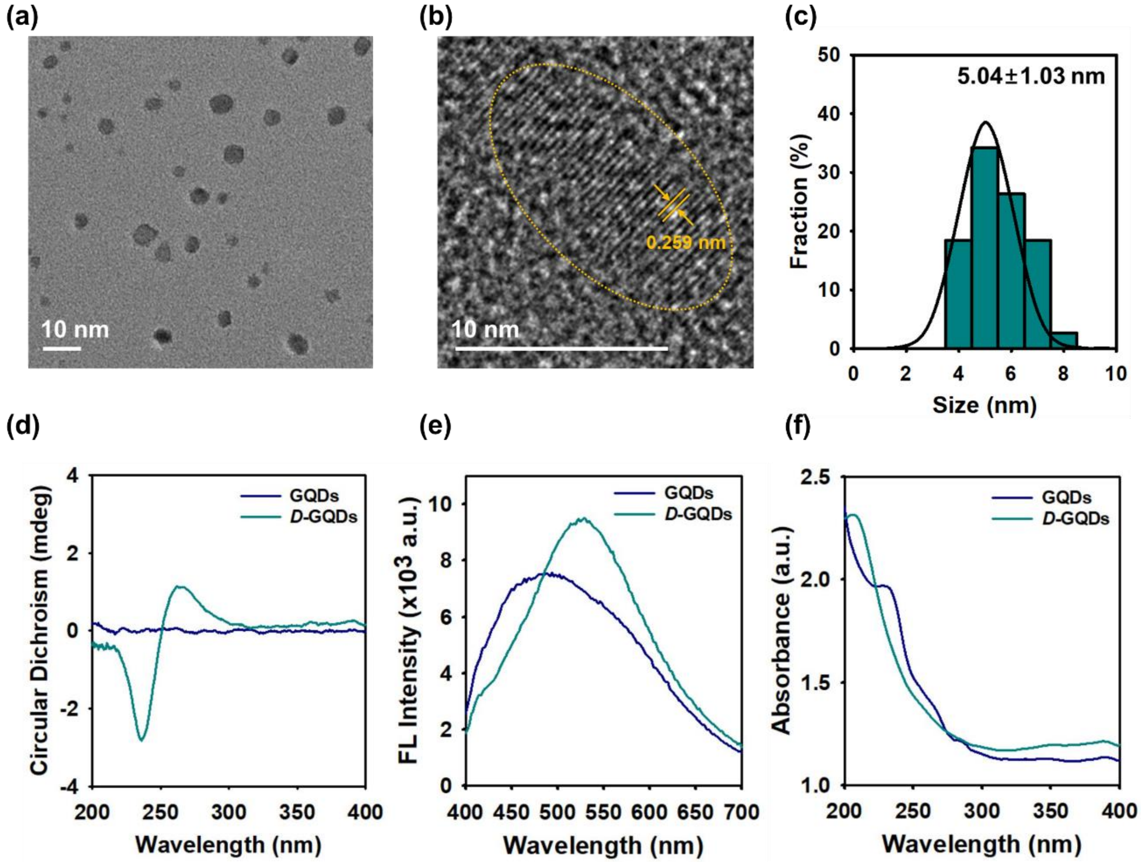


**Figure S2.** Characterizations of GQDs and *D*-GQDs. (a) TEM image of *D*-GQDs. (b) Crystalline grid structure of *D*-GQDs in TEM image. (c) Size distribution of *D*-GQDs based on analysis of TEM images. (d) Circular Dichroism (CD) spectra, (e) fluorescent spectra excited at 360nm, and (f) UV-visible spectra of pristine GQDs and *D*-GQDs.

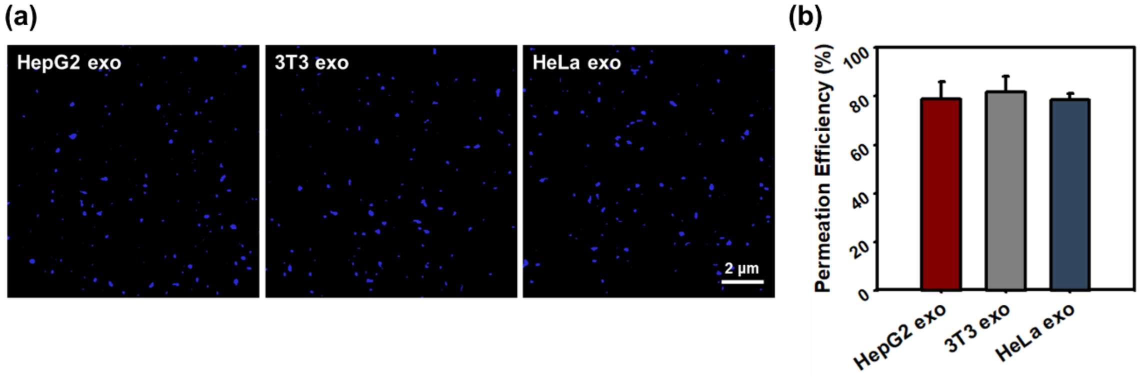


**Figure S3.** Optimal permeation concentration of *D*-GQDs into exosomes. (a) Permeation of *D*-GQDs (blue) into exosomes was observed by CLSM. (b) Permeation efficiency was quantified by counting *D*-GQDs loaded exosomes over the total number of exosomes. The samples were prepared by loading 15 μM *D*-GQDs with exosomes ($1 \times 10^9$ particles/mL).

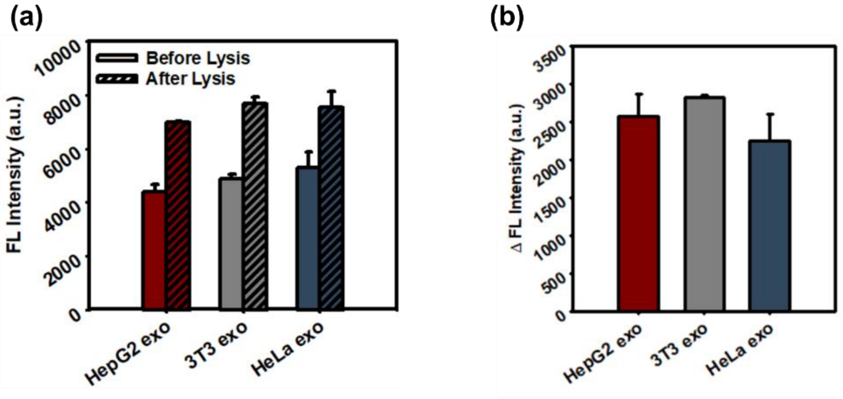


**Figure S4.** (a) Fluorescence intensity before and after lysis of exosome membranes excited at 360nm. (b) Fluorescence recovery of *D*-GQDs loaded exosome lysates. The samples were prepared by loading 12 μM *D*-GQDs with exosomes ($1 \times 10^9$ particles/mL).

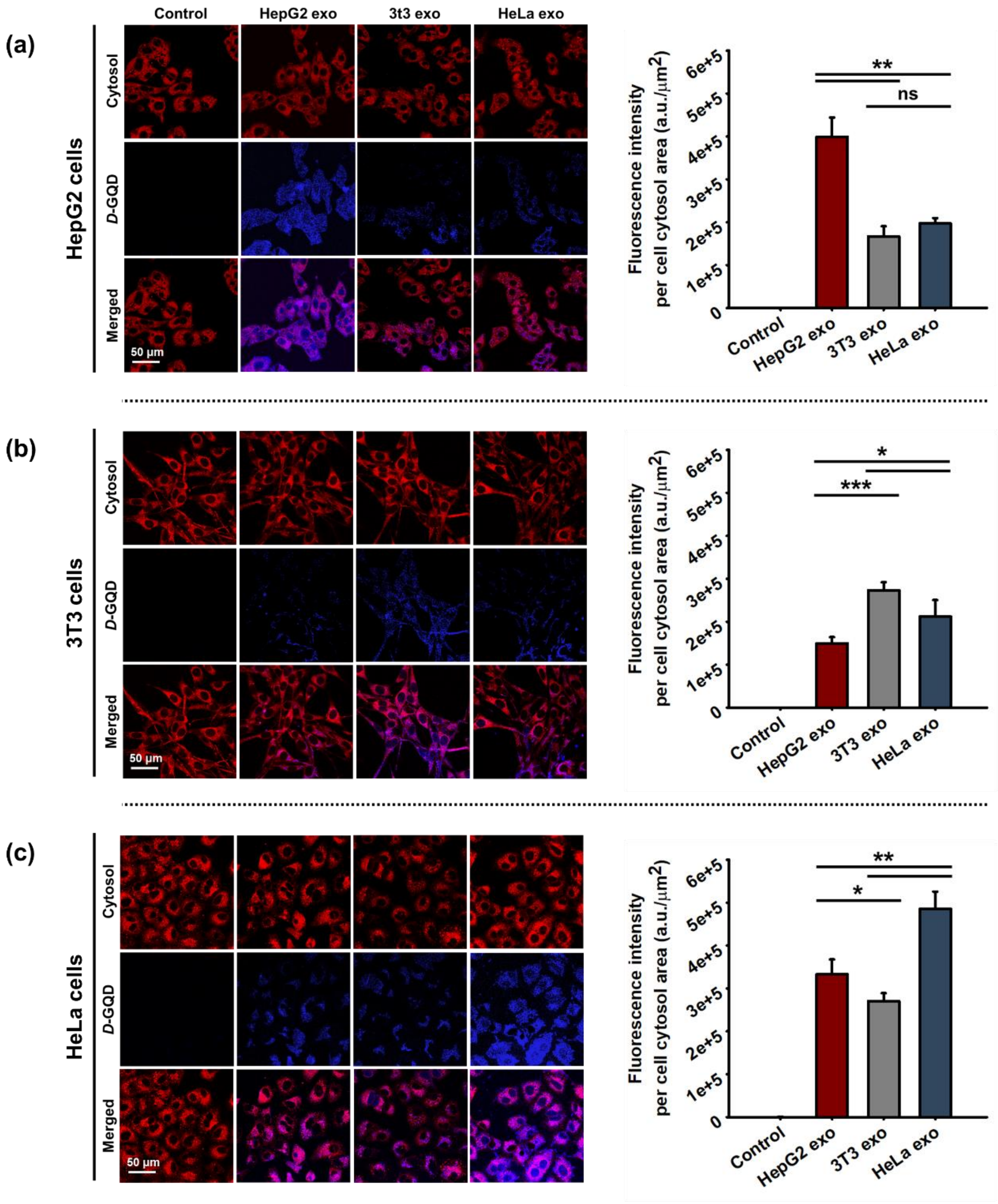


**Figure S5.** Cellular uptake profiles of *D*-GQDs loaded exosomes derived from three different cell lines into (a) HepG2 cells, (b) 3T3 cells, and (c) HeLa cells (mean ± s.d.). ns: not significant, *: $p < 0.05$, **: $p < 0.01$, ***: $p < 0.001$, ****: $p < 0.0001$. Each channel represents: red for the cell cytosol area, and blue for *D*-GQDs. HepG2, 3T3, and HeLa cell lines were incubated for 6 hours with cell culture medium (Control) with a concentration of $0.2 \times 10^3$ exo/cell. The samples were prepared by loading 15 μM *D*-GQDs with exosomes ($1 \times 10^9$ particles/mL).

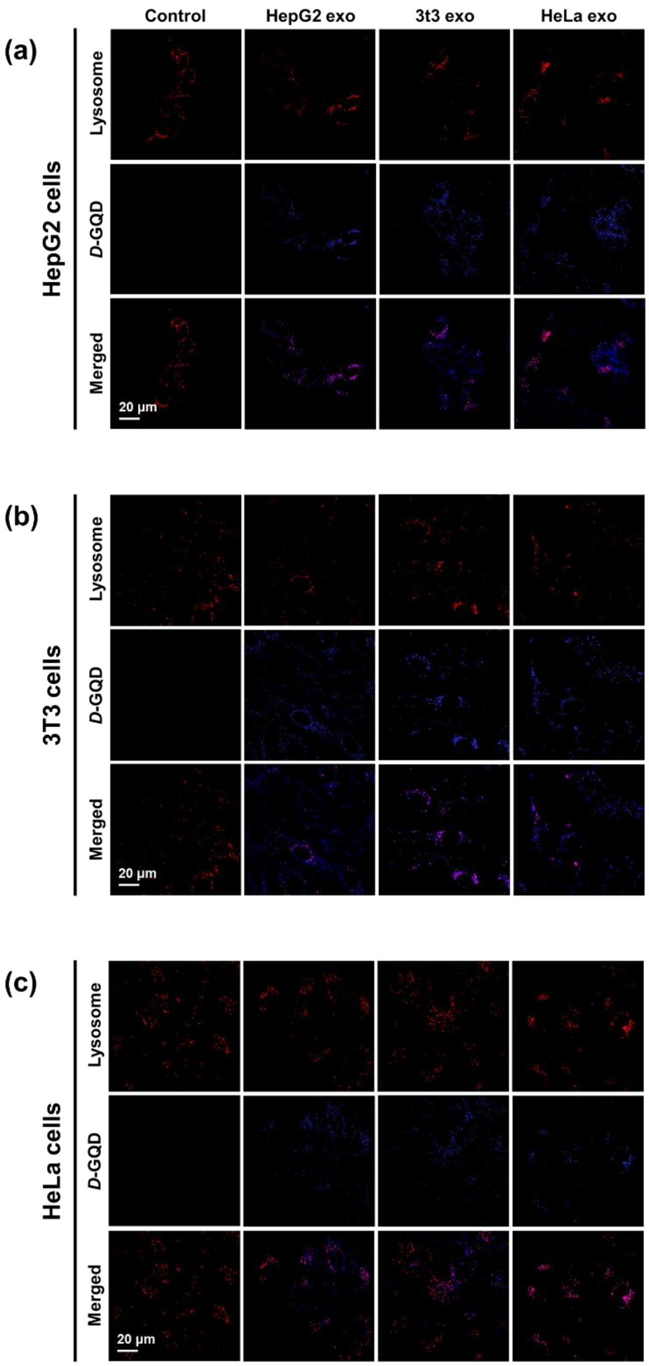


**Figure S6.** Endocytic uptake profiles of exosomes. CLSM images of *D-GQDs* (blue) and lysosomes (red) in (a) HepG2 cells, (b) 3T3 cells, and (c) HeLa cells. Cells were incubated for 4 hours with cell culture medium (Control) with a concentration of 0.2 × $10^3$ exo/cell. The samples were prepared by loading 12 μM *D*-GQDs with exosomes (1 × $10^9$ particles/mL).

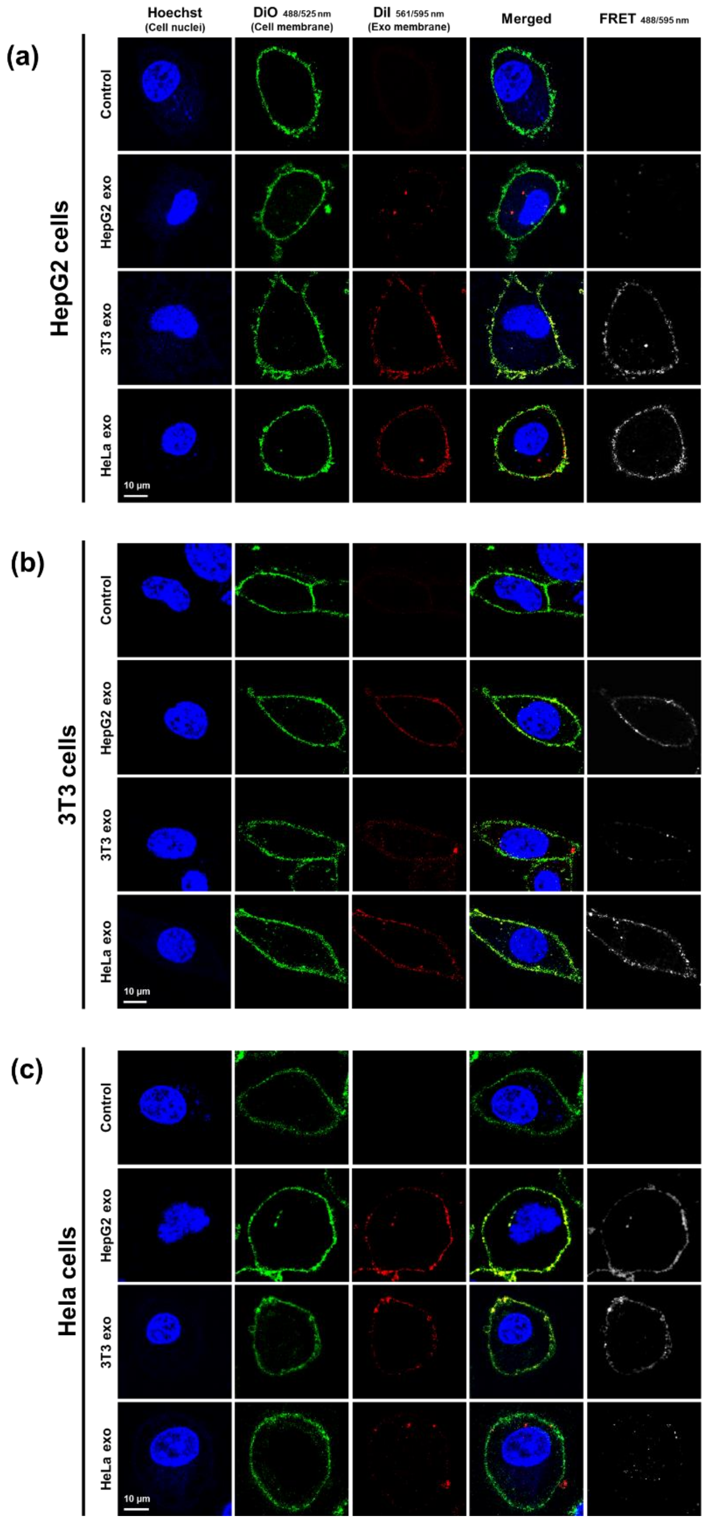


**Figure S7.** Membrane fusion between cells and exosomes. Each channel and merged images of cellular and exosomal membranes was analyzed using CLSM images in (a) HepG2 cells, (b) 3T3 cells, and (c) HeLa cells after 1 hour of incubation with exosomes. Each channel represents: blue for nuclei, green for the cellular membrane, red for the exosomal membrane, and gray for FRET.

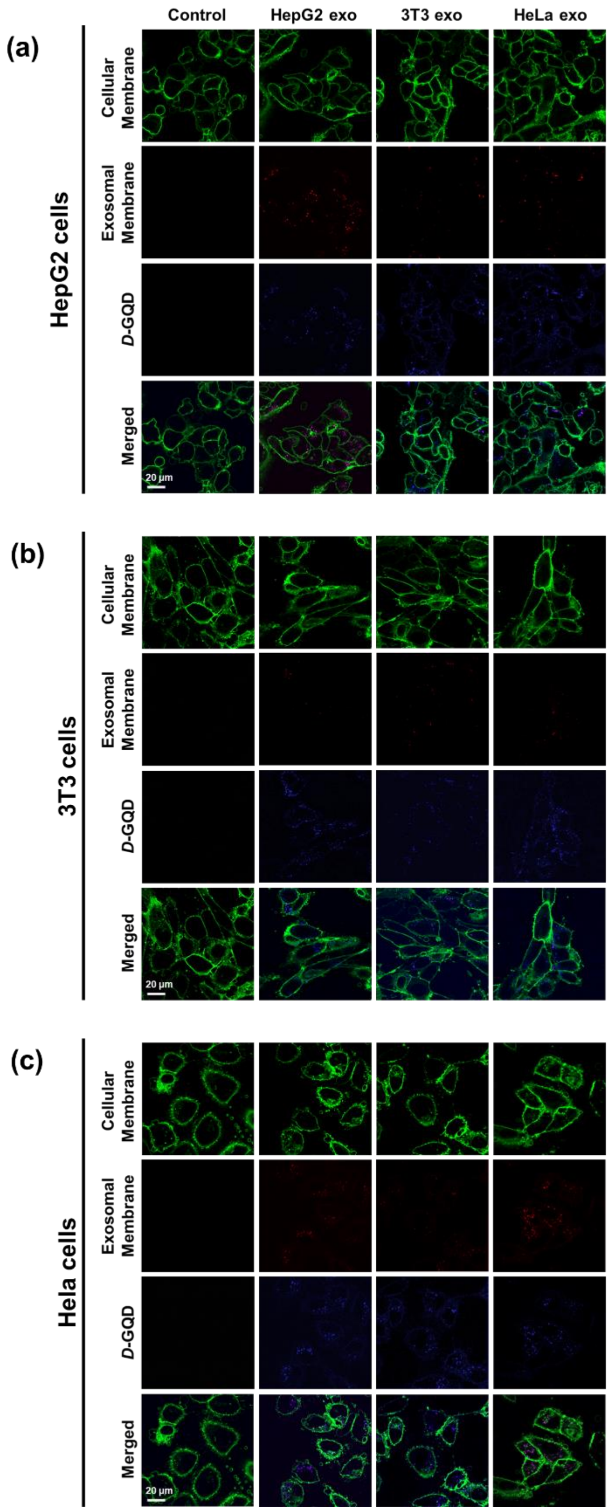


**Figure S8.** *D*-GQDs release and retention from exosomes after 6 hours of incubation with (a) HepG2 cells, (b) 3T3 cells, and (c) HeLa cells. Each channel represents: green for the cellular membrane, red for the exosomal membrane, and blue for *D*-GQDs. Cells were incubated with cell culture medium (Control) with a concentration of $0.2 \times 10^3$ exo/cell. The samples were prepared by loading 12 μM *D*-GQDs with exosomes ($1 \times 10^9$ particles/mL).

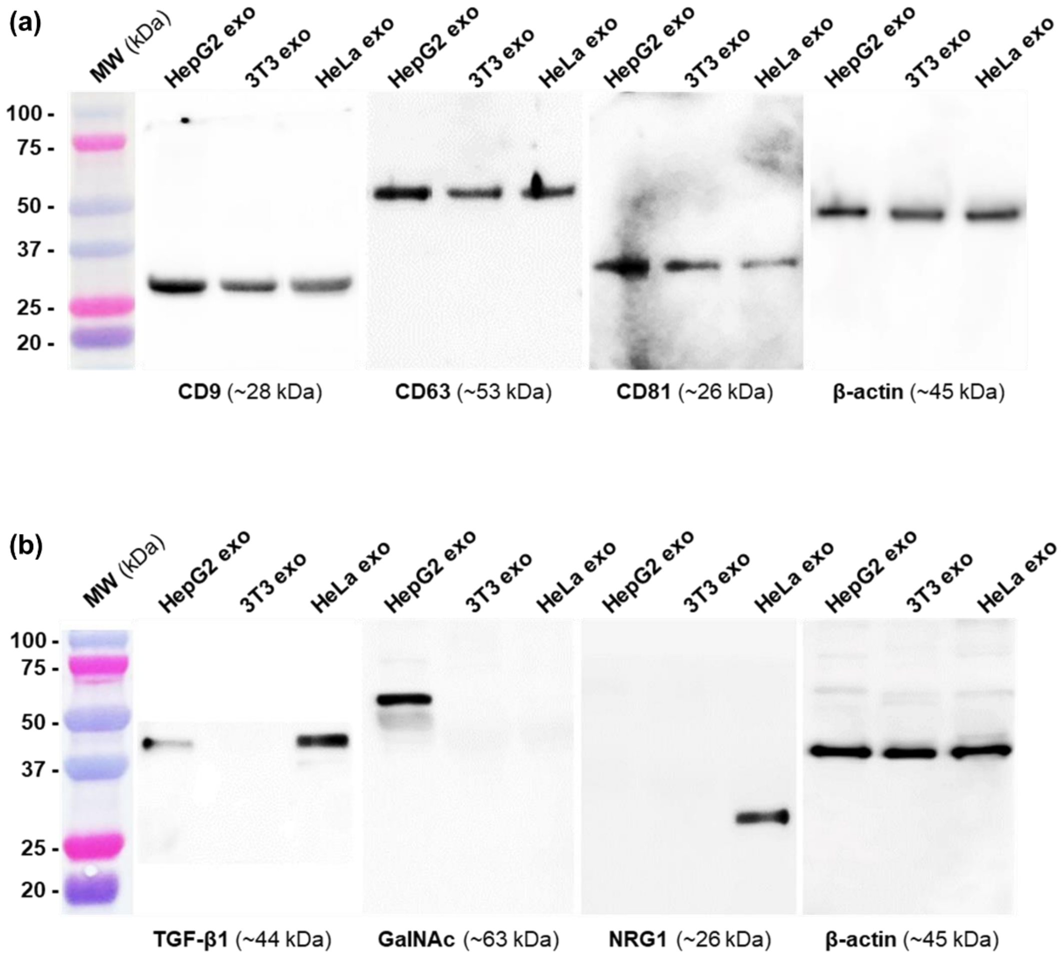


**Figure S9.** Western Blot analysis of (a) exosomal biomarkers (CD9, CD63 CD81) and (b) proteins on exosomes that mediated endocytosis (TGF-β1, GalNAc, NRG1).
